# Single-cell RNA editing defines clinically relevant cellular states in chronic myelomonocytic leukemia

**DOI:** 10.64898/2026.03.15.711339

**Authors:** Nisansala Wickramasinghe, Doan Bui, Surendra Neupane, Meghan C. Ferrall-Fairbanks, Michael M Deininger, Eric Padron, Tongjun Gu

## Abstract

**Background:** Chronic myelomonocytic leukemia (CMML) is a clinically heterogeneous myeloid malignancy with limited therapeutic options and suboptimal risk stratification. Although single-cell RNA sequencing has refined disease classification through gene expression profiling, post-transcriptional mechanisms—particularly adenosine-to-inosine (A-to-I) RNA editing—remain unexplored at single-cell resolution. We hypothesized that cell-specific RNA editing programs contribute to CMML heterogeneity and define distinct, clinically actionable cellular states in CMML.

**Methods:** We developed a single-cell–aware computational framework for high-confidence identification and quantification of RNA editing events. Candidate sites were detected at pseudo-bulk depth using stringent filters and subsequently quantified at single-cell resolution. The pipeline incorporated dual alignment, barcode correction, artifact removal, and exclusion of genomic variants to ensure specificity. We applied this framework to discovery and independent validation CMML cohorts. Editing-defined cellular states were identified by unsupervised clustering of single-cell editing profiles and evaluated for associations with clinical stage, TET2 status, survival, and response to hypomethylating agent (HMA) therapy. Regulatory mechanisms were assessed by analyzing *ADAR1*/*ADAR2* expression and relationships between editing levels and target gene expression.

**Results:** We identified 3,326 high-confidence A-to-I RNA editing sites and delineated reproducible editing-defined cellular states. A granulocyte–monocyte progenitor–like editing state (edClu1_sub0) aligned with an inflammatory, monocytic-biased transcriptional program and was significantly associated with adverse survival, advanced-stage disease and TET2-mutant CMML, supporting it as a high-risk biomarker-defined subpopulation. In contrast, states such as edClu3 and edClu6 were enriched in earlier-stage, TET2–wild-type CMML and correlated with improved outcomes. Editing-defined states demonstrated systematic remodeling following HMA therapy, indicating treatment-responsive post-transcriptional programs. The high-risk state exhibited elevated *ADAR1* and reduced *ADAR2* expression, suggesting enzyme-specific regulatory imbalance as a potential therapeutic vulnerability. Integrative analyses further nominated immune-related genes—including *LAPTM5*, *CTSS*, and *CD83*—as CMML-specific oncogenic RNA editing targets, with coordinated increases in editing and expression within the aggressive state.

**Conclusions:** RNA editing represents a clinically informative and mechanistically relevant layer that refines CMML stratification at single-cell resolution, independent of gene expression. These findings provide a framework for integrating post-transcriptional regulation into precision oncology and highlight RNA editing signatures as biomarkers for risk assessment, treatment monitoring, and therapeutic targeting in hematologic malignancies.

## Background

Chronic myelomonocytic leukemia (CMML) is a clonal hematologic malignancy with overlapping features of myelodysplastic syndromes (MDS) and myeloproliferative neoplasms (MPN)[1, 2]. It exhibits marked clinical heterogeneity, with variable progression risk and therapeutic response. Although recurrent genetic mutations have been well characterized, current molecular stratification incompletely explains clinical variability, and effective treatments remain limited. Allogeneic hematopoietic stem cell transplantation (alloSCT) is the only curative option; however, 5-year overall survival is approximately 20–50%, and only ∼10% of patients are eligible[2]. These limitations highlight the need to identify additional regulatory mechanisms that contribute to CMML pathogenesis and may improve risk stratification and therapeutic targeting.

RNA editing is a post-transcriptional modification that diversifies RNA sequences without altering the DNA genome[3]. In humans, the predominant form is adenosine-to-inosine (A-to-I) editing, catalyzed by the ADAR family of enzymes. Because inosine is interpreted as guanine during reverse transcription and translation, A-to-I editing appears as A-to-G substitutions in RNA sequencing data. By modulating RNA stability, splicing, coding potential, and immune signaling, A-to-I editing regulates key cellular processes and has been increasingly implicated in cancer progression, immune evasion, and therapeutic resistance[4–11]. However, its contribution to CMML biology remains largely unexplored, particularly at single-cell resolution. Whether RNA editing can independently define clinically meaningful cellular states is unknown.

RNA editing detection is technically challenging due to alignment ambiguity at splice junctions, repetitive regions, and among gene isoforms, as well as sequencing errors and amplification artifacts, all of which can lead to high false discovery rates without optimized analytical strategies[12–14]. These challenges are further amplified at single-cell resolution because of sparse coverage and increased susceptibility to technical noise[15]. Consequently, most prior studies rely on bulk measurements or project editing signals onto gene expression–defined clusters, thereby limiting the discovery of post-transcriptionally organized disease states.

To address these limitations, we developed a single-cell–aware computational framework for robust identification and quantification of RNA editing events. Our approach combines stringent pseudo-bulk site discovery with sensitive single-cell quantification, incorporates dual-alignment strategies and barcode-aware duplicate removal, and applies multi-layer artifact filtering to achieve high-confidence editing detection in clinical samples. This framework enables unsupervised clustering based solely on RNA editing profiles, independent of gene expression– based classification.

Applying this approach to CMML scRNA-seq data, we systematically mapped the RNA editing landscape at single-cell resolution and, for the first time, identified RNA editing–defined cellular states that are independent of gene-expression–based clustering. We further characterized their biological, clinical, and mechanistic associations and validated these findings in an independent cohort. Together, these results establish RNA editing as a previously underappreciated organizational layer in CMML and uncover enzyme-linked editing programs associated with disease stage, clinical outcome, and candidate pathways for mechanistic investigation and therapeutic intervention.

## Results

### Overview of the analysis framework

Accurate detection of RNA editing events at single-cell resolution is challenging due to sparse coverage, amplification noise, and alignment ambiguity inherent to scRNA-seq data. To address these limitations, we developed a single-cell aware computational framework that enables robust discovery and quantification of RNA editing events by integrating stringent pseudo-bulk site discovery with sensitive single-cell quantification (**Sup. Fig. 1a**). Candidate RNA editing sites were first identified at the pseudo-bulk level using stringent filtering criteria to leverage higher sequencing depth and minimize false positives. Editing levels at these sites were then quantified in individual cells using relaxed criteria to maximize sensitivity while preserving site-level confidence. This two-phase strategy decouples site discovery from cell-level quantification, enabling reliable detection of RNA editing despite the sparse nature of single-cell data.

**Fig. 1.**
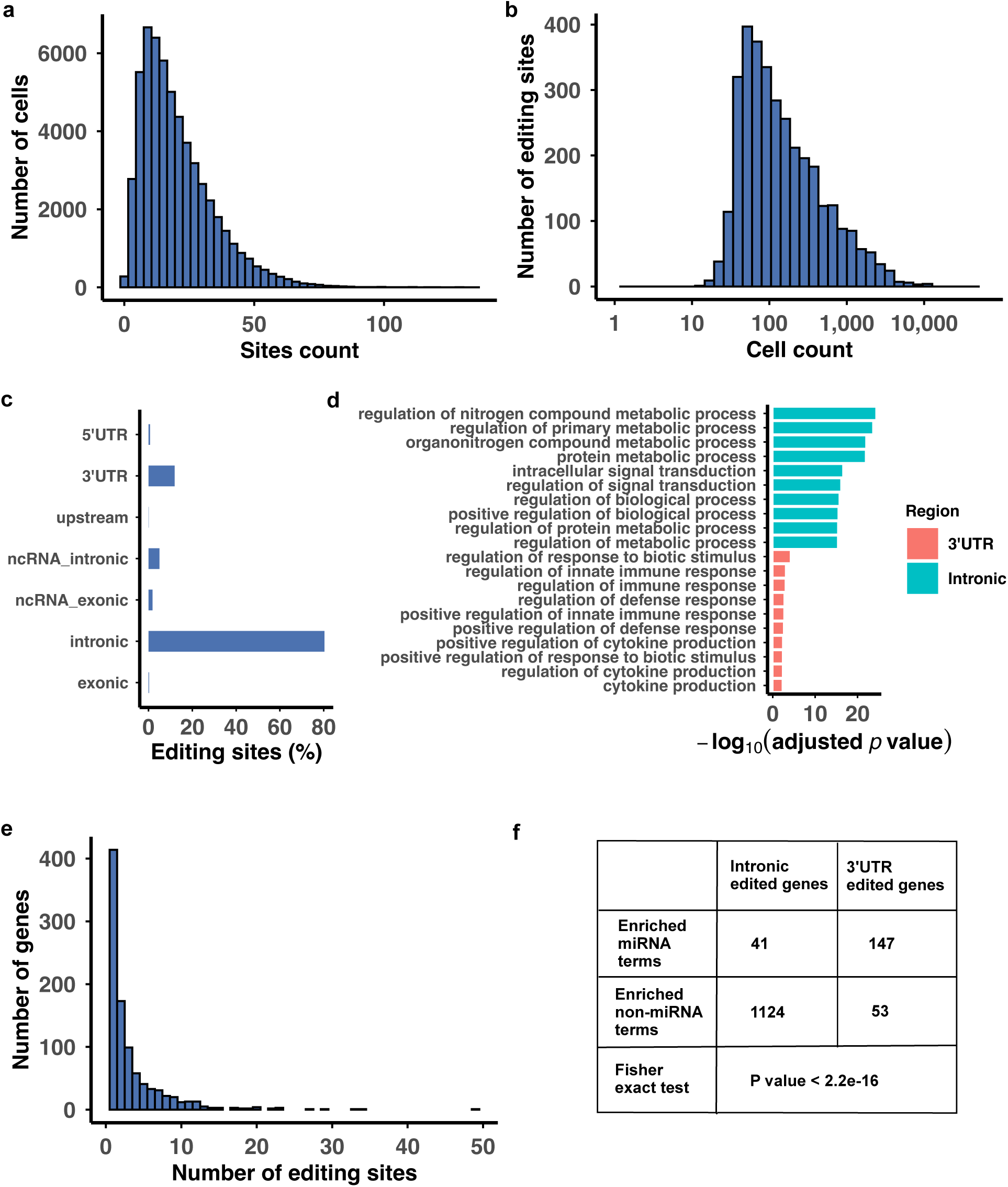
Overview of editing events detected in single cells. (a) Histogram of the number of editing sites per cell. (b) Histogram of the number of single cells per editing site (x axis: log10 scale). (c) Percentage of editing sites in different genomic regions. (d) Top 10 enriched GO:BP functions for genes harboring intronic and 3’UTR editing sites. (e) Distribution of the number of editing sites located in genes. (f) 3’UTR edited genes significantly enriched miRNA terms compared to intronic edited genes.

Unlike most prior RNA editing studies that define cellular clusters based on gene expression and subsequently project editing signals onto those clusters, our framework directly quantifies RNA editing at single-cell resolution, independent of transcriptional clustering. To improve cell barcode fidelity, sequenced barcode sequences were compared with reference barcode sets and corrected when mismatches fell within a tolerated threshold, increasing retention of usable reads while maintaining accurate cell assignment. Read alignment was performed using a dual-aligner strategy combining Bowtie[16] and STAR[17], including dual-reference alignment for Bowtie, to enhance alignment sensitivity and fidelity. Potential PCR duplicates were removed on a per-cell basis rather than at the sample level, retaining more usable reads for downstream single-cell analyses.

Multiple post-alignment filters were applied to minimize technical and genomic artifacts, including strand-bias filtering using Fisher’s exact test, positional bias filtering based on distance to read ends, exclusion of sites located in simple repeat regions, and removal of sites overlapping known genomic variants. We retained only RNA editing sites with an editing ratio greater than 0.05 in at least 10 patients, yielding a high-confidence set of recurrent RNA editing events suitable for downstream analyses. Full details of the computational pipeline are provided in the Methods.

A total of 24 CMML patients were used as the discovery cohort. An independent validation cohort of 13 CMML patients was reserved exclusively for validation and was not used at any stage of the discovery process, ensuring unbiased assessment of reproducibility [18]. Sample and patient characteristics are summarized in **Sup. Tab. 1**. Three healthy individuals from the study of Hua *et al*.[19] as references for identifying healthy-like cells (**Sup. Tab. 1**).

**Tab. 1.** GMP-specific oncogenic editing sites.

| Gene | Number of editing sites/Gene | Editing sites (chr:position:strand) |
| --- | --- | --- |
| ACSL1 | 1 | 4:184822487:- |
| ADAMTSL4-AS1 | 1 | 1:150565493:- |
| ADGRE5 | 1 | 19:14398554: + |
| AGFG1 | 1 | 2:227539541:+ |
| AKAP13 | 1 | 15:85587883:+ |
| ANKRD11 | 1 | 16:89337630:- |
| ARPC2 | 3 | 2:218235855:+, 2:218235856:+, 2:218235983:+ |
| ARRB2 | 1 | 17:4713551:+ |
| BACH1 | 1 | 21:29337849:+ |
| CD83 | 5 | 6:14121360:+, 6:14121438:+, 6:14121439:+, 6:14121525:+, 6:14121547:+ |
| CSGALNACT2 | 1 | 10:43141467:+ |
| CTDSP1 | 1 | 2:218403975:+ |
| CTSS | 2 | 1:150732772:-, 1:150732802:- |
| CYRIB | 4 | 8:130001625:-, 8:130001667:-, 8:130001747:-, 8:130001752:- |
| FBXO34 | 1 | 14:55279055:+ |
| GAS7 | 1 | 17:10133242:- |
| GPCPD1 | 2 | 20:5573178:-, 20:5601569:- |
| HAVCR2 | 1 | 5:157090250:- |
| HCK | 1 | 20:32094613:+ |
| ICAM1 | 1 | 19:10276047:+ |
| IQSEC1 | 2 | 3:13001108:-, 3:13001227:- |
| ITGAX | 1 | 16:31364784:+ |
| LAPTM5 | 3 | 1:30743975:-, 1:30743982:-, 1:30743984:- |
| LINC01619 | 1 | 12:92127570:- |
| LITAF | 3 | 16:11565150:-, 16:11565220:-, 16:11575983:- |
| LYN | 11 | 8:55906537:+, 8:55906540:+, 8:55906554:+, 8:55906691:+, 8:55906694:+, 8:55922682:+, 8:55922687:+, 8:55922717:+, 8:55987251:+, 8:55987256:+, 8:55987260:+ |
| MTO1 | 1 | 6:73506903:+ |
| NEAT1 | 4 | 11:65441427:+, 11:65442619:+, 11:65442678:+, 11:65442701:+ |
| PGS1 | 1 | 17:78422912:+ |
| PIK3R5 | 2 | 17:8943374:-, 17:8963658:- |
| PLAUR | 1 | 19:43660408:- |
| PLXDC2 | 3 | 10:19838928:+, 10:19852345:+, 10:19876541:+ |
| RAB7A | 1 | 3:128744060:+ |
| RRP12 | 3 | 10:97377018:-, 10:97377020:-, 10:97386477:- |
| SH3BP2 | 3 | 4:2840110:+, 4:2840114:+, 4:2840118:+ |
| SIPA1L1 | 6 | 14:71356515:+, 14:71440563:+, 14:71453803:+, 14:71453808:+, 14:71453847:+, 14:71453929:+ |
| SLC2A3 | 2 | 12:7923842:-, 12:7923975:- |
| SLC6A6 | 1 | 3:14433673:+ |
| SPI1 | 1 | 11:47367896:- |
| STK10 | 1 | 5:172112823:- |
| TEX14 | 1 | 17:58624479:- |
| TGFB1 | 1 | 19:41341277:- |
| ZFAND3 | 2 | 6:37833679:+, 6:37884339:+ |
| ZSWIM6 | 6 | 5:61365254:+, 5:61365258:+, 5:61378938:+, 5:61379058:+,<br>5:61379060:+, 5:61379106:+ |

### Single-cell RNA editing landscape in CMML

Across the 24 CMML samples, we obtained an average of 2,495 cells per patient and an average of ∼125,009 reads per cell after filtering the cells with less than 50,000 reads, with a mean unique alignment rate of ∼61% per sample. At the pseudo-bulk level, we identified 509,539 A-to-G and four C-to-U RNA editing sites. All A-to-G sites were registered in REDIportal entries, a curated RNA editing database, supporting high confidence of our site discovery pipeline. The distribution of all identified variant sites, including potential DNA mutations, is shown in **Sup. Fig. 1b**. A-to-G alteration predominated with non-A-to-G variants accounting for less than 8.3% of all detected sites.

At the single-cell resolution, we detected 3,326 editing sites across 56,940 CMML cells. Most RNA editing sites were present in only a subset of cells, and most cells harbored only a small fraction of these sites. Yet a subset of RNA editing sites occurs in thousands of cells (**Fig. 1a-1b**), indicating heterogeneous but structured editing activity across the CMML cellular landscape. Annotation of these editing sites identified 961 host genes (**Sup. Tab. 2**). Gene Ontology (GO) analysis of host genes revealed enrichment for metabolic processes, stress responses, and intracellular signaling pathways (**Sup. Tab. 3**). Most editing events were located within intronic region and the 3′ untranslated region (3′UTR) (**Fig. 1c**), consistent with previous studies in other cancers[6, 10, 20]. While intronic sites mirrored overall GO enrichments, genes harboring 3’UTR editing sites were specifically enriched for innate immune response and cytokine production pathways (**Fig. 1d**; **Sup. Tab. 4-5**). A small number of editing sites were also located within miRNA genes or their host transcripts (**Sup. Tab. 6**). Although most genes harbored one or two editing sites, 70 genes contained ≥10 editing sites (**Fig. 1e**). These highly edited genes were enriched for functions related to leukocyte-mediated immunity and regulation of signaling, consistent with CMML biology (**Sup. Tab. 7**). Notably, genes with 3′UTR editing sites showed significantly stronger enrichment for miRNA-related terms compared with genes harboring intronic editing sites (**Fig. 1f; Sup. Tab. 4-5**).

Together, these analyses define a structured single-cell RNA editing landscape in CMML characterized by heterogeneous cellular distribution, distinct genomic localization patterns, and functionally coherent enrichment signatures.

### RNA editing defines discrete cellular states independent of gene expression

To determine whether RNA editing patterns alone can define cellular organization, we performed unsupervised clustering of single cells using RNA editing profiles without incorporating gene expression information. Using editing ratios across the 3,326 high-confidence RNA editing sites, we identified nine major RNA editing-defined clusters (edClu0–edClu8), with additional substructures observed within edClu1 (**Fig. 2a**). Except for edClu7 and edClu8, which contained relatively few cells and marker sites, the remaining clusters comprised substantial numbers of cells and exhibited robust, cluster-specific RNA editing profiles (**Sup. Fig. 2a; Sup. Tab. 8**). Manual inspection further confirmed that top marker editing sites were highly enriched within their corresponding clusters, supporting the specificity of the RNA editing-defined clustering (**Sup. Fig. 2b-2h**).

**Fig. 2.**
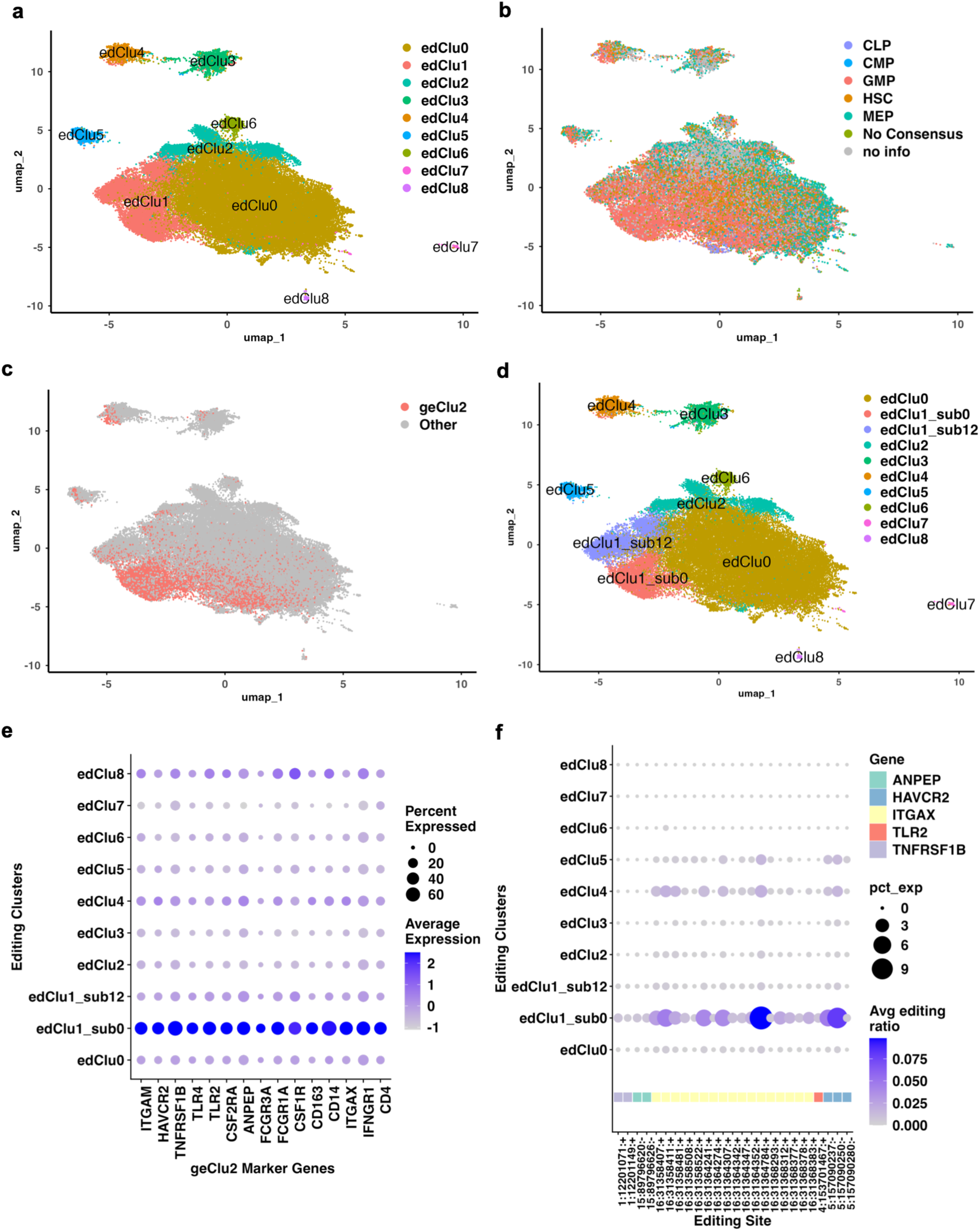
Identification of RNA editing-defined clusters. (a) RNA editing-defined clusters identified based on single-cell RNA editing profiles. (b) Editing cells annotated with gene expression-based cell type identities. (c) Cells assigned to the aggressive gene expression cluster, geClu2. (d) RNA editing-defined clusters following subclustering of edClu1. (e) Scaled and normalized average expression and percentage of cells expressing geClu2 marker genes across RNA editing-defined clusters. (f) Average RNA editing ratios for editing sites located within geClu2 marker genes across RNA editing-defined clusters.

To evaluate whether RNA editing-defined clusters correspond to biologically and clinically meaningful cellular states, we leveraged previously established gene expression–based classification from the same CMML samples, as reported by our coauthors Ferrall-Fairbanks *et al*.[18]. That study identified a clinically aggressive granulocyte–monocyte progenitor (GMP)–like cluster, geClu2, a subset of the GMP cell type associated with poor prognosis. By transferring both cell-type annotations and gene expression cluster labels from that study onto our RNA editing-defined cells (**Fig. 2b**), we observed that edClu1 was predominantly composed of GMP-like cells. Notably, a subregion of edClu1 showed substantial overlap with the adverse geClu2 transcriptional cluster (**Fig. 2c**).

Further sub-clustering analysis of edClu1 revealed three subpopulations, edClu1_sub0, edClu1_sub1, and edClu1_sub2, among which geClu2 cells were most strongly enriched in edClu1_sub0 (**Fig. 2d**). Accordingly, we merged edClu1_sub1 and edClu1_sub2 into a single group (edClu1_sub12) and focused subsequent analyses on edClu1_sub0. Marker genes defining geClu2 were significantly enriched in edClu1_sub0 (**Fig. 2e**) and RNA editing events located within these marker genes were also predominantly observed in edClu1_sub0 (**Fig. 2f**). These findings demonstrate that RNA editing–based clustering, derived independently of gene expression, recapitulates a clinically aggressive GMP-associated cellular state previously defined by gene expression profiling in CMML.

### Lineage-associated RNA editing programs underlie editing-defined states

To elucidate the biological basis of RNA editing-defined cellular states, we examined the distribution of annotated hematopoietic cell types across editing-defined clusters. EdClu1_sub0 and edClu1_sub12 were predominantly composed of GMP cells, whereas edClu2, edClu3, edClu5, edClu6, and edClu7 were enriched for megakaryocyte–erythroid progenitor (MEP) cells (**Sup. Fig. 2i**).

To quantify lineage-associated RNA editing activity, we computed cell-type–specific mean editing levels (MELs) by averaging editing ratios across lineage-specific marker editing sites. GMP-associated marker sites exhibited the highest MELs in edClu1_sub0 (**Fig. 3a–b**), whereas MEP-associated markers were most elevated in edClu2, edClu3, edClu5, and edClu6 (**Fig. 3c–d**). These results confirm that edClu1_sub0 represents a GMP-dominated editing state and demonstrate that multiple MEP-enriched clusters share elevated lineage-specific RNA editing activity. A complete list of lineage-specific marker editing sites is provided in **Sup. Tab. 9**.

**Fig. 3.**
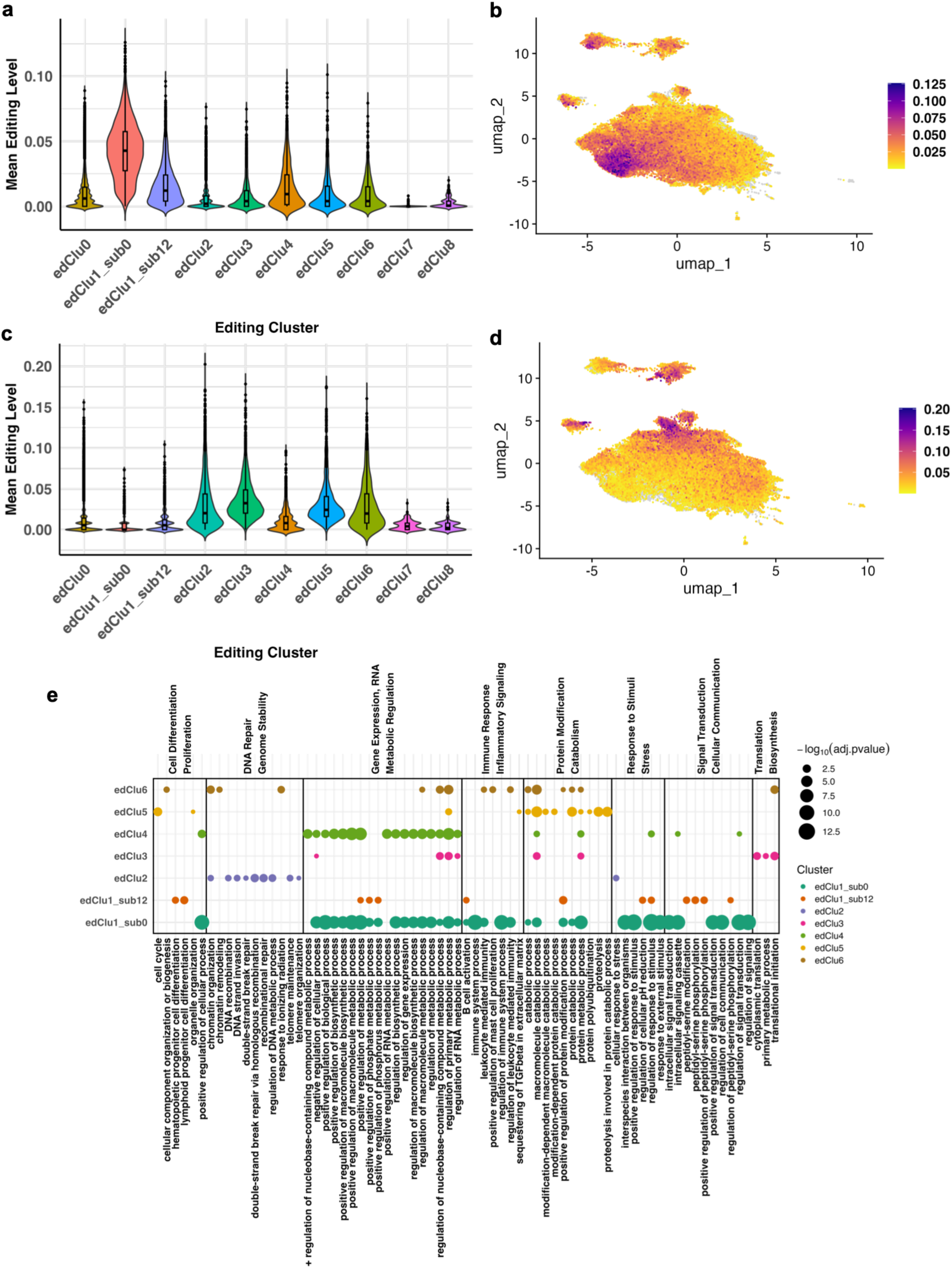
Cell-type–specific RNA editing patterns and functional enrichment across editing clusters. (a, b) Mean editing levels of GMP-associated marker sites across editing-defined clusters. (c, d) Mean editing levels of MEP-specific marker sites across editing-defined clusters. (e) Gene Ontology Biological Processes associated with genes harboring marker editing sites of each cluster. Dot size represents –log₁₀ adjusted p-value, reflecting the significance of enrichment. Here, the top 15 biological process terms per cluster were selected, grouped and visualized based on functional similarity.

We next performed functional enrichment analysis of genes harboring cluster-specific RNA editing sites to further characterize the biological programs associated with each editing-defined state (**Fig. 3e; Sup. Tab. 10–11**). EdClu1_sub0 was enriched for pathways related to inflammation, immune response, and stress and stimulus signal transduction, consistent with previously described inflammatory GMP-like programs in CMML associated with advanced disease features and poor clinical outcomes(14). EdClu2 showed enrichment for pathways involved in DNA recombination and repair, telomere maintenance, and homologous recombination, suggesting enhanced genomic maintenance and cellular stability. In contrast, clusters edClu3–6 exhibited shared enrichment for metabolic pathways, including common processes such as RNA metabolic regulation and protein modification and catabolism, alongside cluster-specific signatures. For example, edClu4 showed stronger enrichment for macromolecule biosynthetic and metabolic processes. A curated summary of the top enriched GO biological process terms for each cluster is shown in **Sup. Fig. 2j**. These analyses indicate that RNA editing-defined states reflect structured, lineage-associated regulatory programs rather than arbitrary cellular groupings.

### RNA editing-defined states associate with treatment response and disease stage

To assess whether RNA editing-defined cellular states are associated with clinical characteristics in CMML, we examined the distribution of cells within editing-defined clusters across clinical subgroups and treatment conditions. For each sample, we calculated both the proportion of the cells in each editing-defined cluster out of all cells assigned to all clusters and MELs, then compared across clinical contexts.

We first evaluated changes in cluster composition following hypomethylating agent (HMA) therapy using five treated and untreated normal-like patients, as defined by Ferrall-Fairbanks *et al.*[18]. Following HMA treatment, clusters edClu2, edClu3, edClu4, edClu5, and edClu6 exhibited marked increases in both cellular proportions and MELs (**Fig. 4a-4b**), indicating that RNA editing activity within these clusters may play protective role in normal-like CMML patients.

**Fig. 4.**
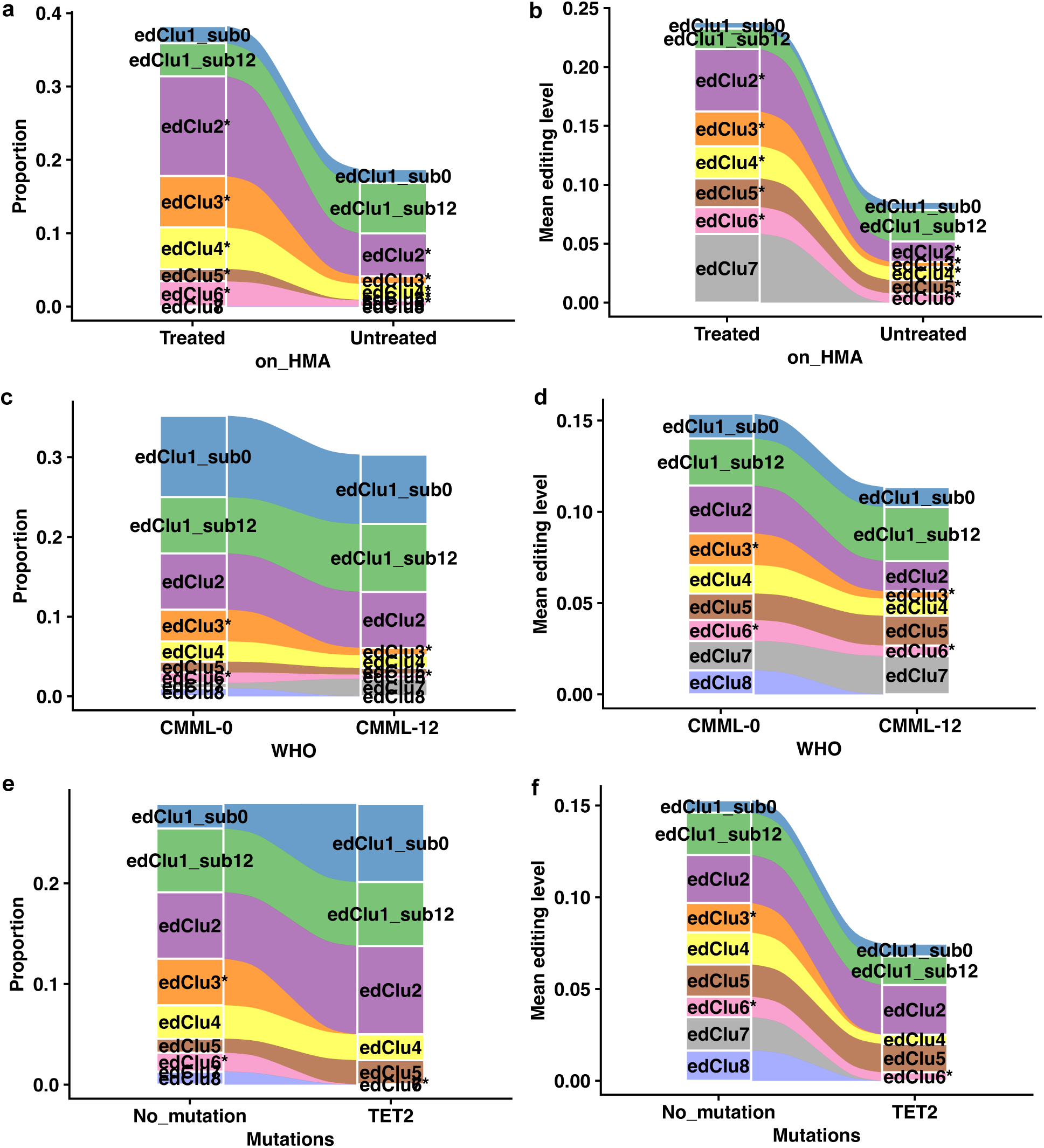
Alterations in RNA editing-defined clusters across clinical features. (a,c,e) Association between clinical features and the proportion of cells across editing-defined clusters. For each sample, the proportion of cells assigned to each editing cluster was calculated as the number of cells in the cluster divided by the total number of cells in the sample. (b,d,f) Association between mean editing level (MEL) and clinical features. MEL was calculated for each editing site by averaging the editing ratios of all cells within each editing cluster and sample. These values were then averaged across all marker editing sites for each cluster. * denotes clusters exhibiting apparent changes.

To assess associations with disease progression, we compared cluster cellular proportion and MELs across WHO CMML subtypes using untreated samples to avoid treatment-related confounding. EdClu3 and edClu6 were more abundant in the CMML-0 compared with more advanced CMML-1/2 cases (combined as CMML-12) (**Fig. 4c-4d**). Logistic regression models using cluster-specific MELs as predictors further demonstrated that edClu2 and edClu3 significantly discriminated CMML-0 from CMML-12, with likelihood ratio test P values of 0.0175 and 0.0342 and corresponding areas under the receiver operating characteristic curve (AUCs) of 0.778 and 0.722, respectively (**Sup. Fig. 2k**). Stratification by *TET2* mutation status further revealed that edClu3 and edClu6 exhibited lower proportions and MELs in *TET2*-mutant patients compared with those without *TET2* mutations cases (**Fig. 4e-4f**).

Collectively, these analyses demonstrate that some RNA editing-defined states—particularly edClu3 and, to a lesser extent, edClu6—are preferentially associated with less advanced disease stages and exhibit dynamic modulation in response to epigenetic therapy.

### Prognostic relevance of RNA editing-defined cellular states

To evaluate the prognostic significance of RNA editing-defined cellular states in CMML, we performed survival analyses using Cox proportional hazards models based on cluster cellular proportions and MELs. All models were adjusted for age and sex, and analyses were restricted to patients who had not received HMA therapy to minimize treatment-related confounding.

When modeling cluster cellular proportions, significant survival associations were identified for edClu3, edClu7, and edClu1_sub12 (**Sup. Tab. 12**). In parallel analyses using cluster-specific MELs as predictors, edClu3, edClu6, and edClu1_sub12 showed significant associations with overall survival (**Sup. Tab. 13**). Among these, edClu3 was consistently associated with survival across all models, identifying it as the most robust prognostically relevant RNA editing-defined state.

Consistent with its prognostic associations, edClu3 also exhibited concordant patterns across disease-stage and treatment-response analyses, including enrichment in less advanced CMML subtypes, differential abundance by *TET2* mutation status, and increased RNA editing activity following HMA treatment in normal-like patients (see **the previous section**). EdClu6 emerged as a secondary but consistent signal, showing significant survival associations in MEL-based models and similar directional trends across clinical analyses (see **the previous section**). Taken together, these results identify edClu3 as the most robust prognostically informative RNA editing-defined state in CMML, with edClu6 showing a secondary but concordant association.

### Identification of CMML-associated oncogenic RNA editing programs

To identify RNA editing events associated with malignant hematopoiesis in CMML, we distinguished cancer-like and healthy-like cells at single-cell resolution using inferred copy number variation (CNV) profiles. Using inferCNV with healthy donors as references, cells from the 24 CMML patients were classified into cancer-like and healthy-like populations (see **Methods**), yielding 7,873 cancer-like and 6,325 healthy-like cells after integration with the RNA editing dataset.

We performed multiple complementary validation analyses, detailed in the **Supplementary Materials**, to support the high confidence of these classifications. These included a simulation study, comparisons between cells from healthy donors and inferred healthy-like cells, comparisons between hematopoietic stem cells (HSCs) and healthy-like cells, concordance between geClu2 cells and cancer-like cells, enrichment of cancer-like cells within GMP populations, and evaluation of CMML CD34⁺ marker gene expression in cancer-like versus healthy-like cells (**Sup. Fig. 3a-3k**). Across all validation strategies, we observed consistent and concordant results, supporting the robustness of the inferred malignant and non-malignant cell assignments.

Differential RNA editing analysis between cancer-like and healthy-like cells identified 167 editing sites significantly enriched in cancer-like cells and 109 sites enriched in healthy-like cells (**Sup. Tab. 14**). Genes harboring cancer-enriched editing sites were predominantly associated with immune-related and defense-response processes, whereas healthy-like–enriched sites were linked to RNA metabolic signaling pathways (**Sup. Fig. 3l; Sup. Tab. 15**). In addition, 428 editing sites were detected exclusively in cancer-like cells and 70 were unique to healthy-like cells (**Sup. Tab. 16**). Functional enrichment of these cell-type specific sites recapitulated the same immune and inflammatory themes in cancer-like cells and highlighted both metabolic and intracellular signaling pathways in healthy-like cells (**Sup. Fig. 3m; Sup. Tab. 17**). Notably, these malignancy-associated RNA editing patterns were concordant with RNA editing-defined cellular states. The clinically aggressive GMP-associated cluster, edClu1_sub0, was enriched for inflammatory and defense-response pathways, whereas edClu3 and edClu6, MEP–associated clusters linked to less advanced CMML, showed enrichment for RNA and protein metabolic pathways.

To further link RNA editing with CMML disease features, we performed binomial regression analyses to identify editing sites associated with monocyte burden, adjusting for age, sex, and ADAR1 and ADAR2 expression. This analysis identified 419 editing sites positively and 607 negatively associated with absolute monocyte count, as well as 382 positively and 620 negatively associated sites for monocyte percentage (**Sup. Tab. 18-19**).

Finally, we integrated multiple lines of evidence to prioritize high-confidence oncogenic RNA editing sites. Sites were selected based on: 1) positive association with monocyte absolute count or percentage; 2) enrichment in edClu1_sub0, the most clinically aggressive GMP-associated editing state; 3) cancer-specific editing sites or sites with significant upregulation in cancer-like cells. This intersection yielded 92 high-confidence GMP-specific oncogenic editing sites (oncoEdits_GMP) mapping to 44 genes (**Tab. 1**; **Fig. 5a**). These genes were enriched for immune-related biological processes, including leukocyte-mediated cytotoxicity, immune effector process, MAPK cascade pathway, and cell-cell adhesion (**Sup. Tab. 20**; **Fig. 5b**).

**Fig. 5.**
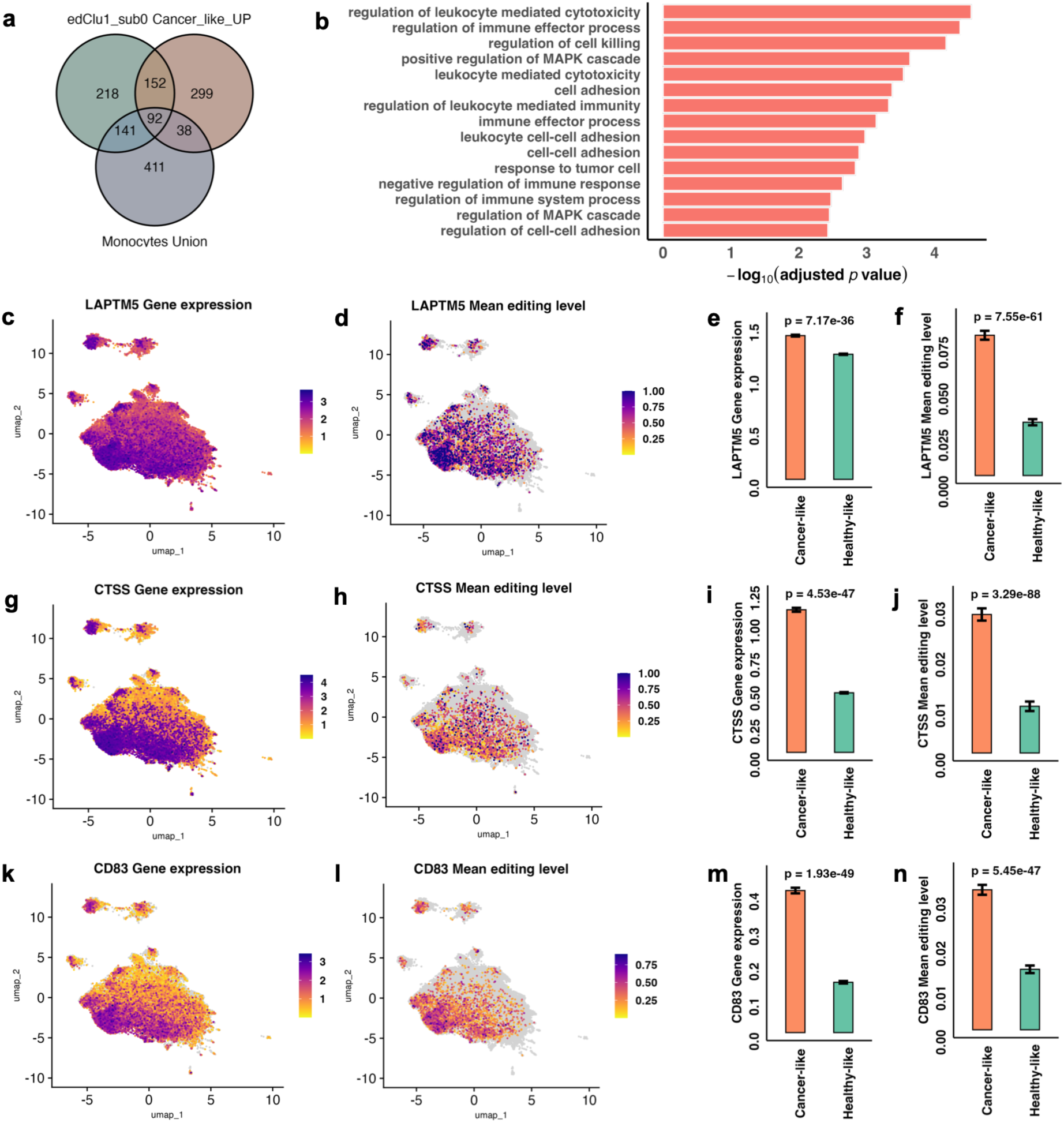
Identification of high confident oncogenic editing sites. (a) Identification of high-confidence, GMP-specific oncogenic editing sites. (b) Enriched functions for the genes with high-confidence, GMP-specific oncogenic editing sites. (c,j,k) Gene expression profile of the genes of interest. (d,h,l) Mean editing levels of the genes of interest. (e,i,m) Differential expression of the genes of interest between healthy-like and cancer-like cells. (f,j,n) Differential mean editing levels of the genes of interest between healthy-like and cancer-like cells.

Several genes—notably *LAPTM5*, *CTSS*, and *CD83*—harbored multiple editing sites, highlighting them as potential key drivers of CMML-associated immune dysregulation. Among these, *LAPTM5* exhibited markedly elevated expression and increased RNA editing within edClu1_sub0 (**Fig. 5c-5f**). Given its established roles in immune regulation and tumorigenesis across multiple solid tumors[21–24], the coordinated overexpression and RNA editing of *LAPTM5* suggest a functional link between RNA editing and immune-related gene regulation in CMML. Similar coordinated expression and editing patterns were observed for *CTSS*[25, 26] (**Fig. 5g-5j**) and *CD83*[27, 28] (**Fig. 5k-5n**), both of which have established roles in immune modulation and tumor biology. Although these genes have not previously been implicated in CMML, their cluster-specific RNA editing and expression patterns support a convergent role for RNA editing in shaping immune-associated oncogenic programs in aggressive CMML.

### Reproducibility of RNA editing-defined states and oncogenic programs in an independent cohort

To evaluate the robustness of RNA editing-defined cellular states, we analyzed an independent cohort of 16 CMML samples, of which 13 samples with sufficient numbers of detected RNA editing sites were retained for downstream analyses (**Sup. Tab. 1**). The validation cohort was processed using the same computational pipeline and parameters as the discovery cohort. Compared with the discovery cohort, which included seven mono-biased patients (∼30%), the validation cohort contained only two mono-biased patients (∼15%), resulting in fewer cells associated with aggressive monocytic programs.

Across 25,110 cells from the 13 validation samples, we identified 1,308 RNA editing sites and performed unsupervised clustering based solely on RNA editing profiles, revealing eight RNA editing-defined clusters (valClu0–valClu7) (**Fig. 6a**). Examination of gene expression derived cell-type annotations showed that GMP cells were predominantly enriched in valClu2 and valClu5 (**Fig. 6b**). HSCs, less distinctly separated, were distributed across valClu0, valClu3, and valClu6 (**Sup. Fig. 4a**). MEP cells were mainly localized in valClu1 and valClu4 (**Sup. Fig. 4b**).

**Fig. 6.**
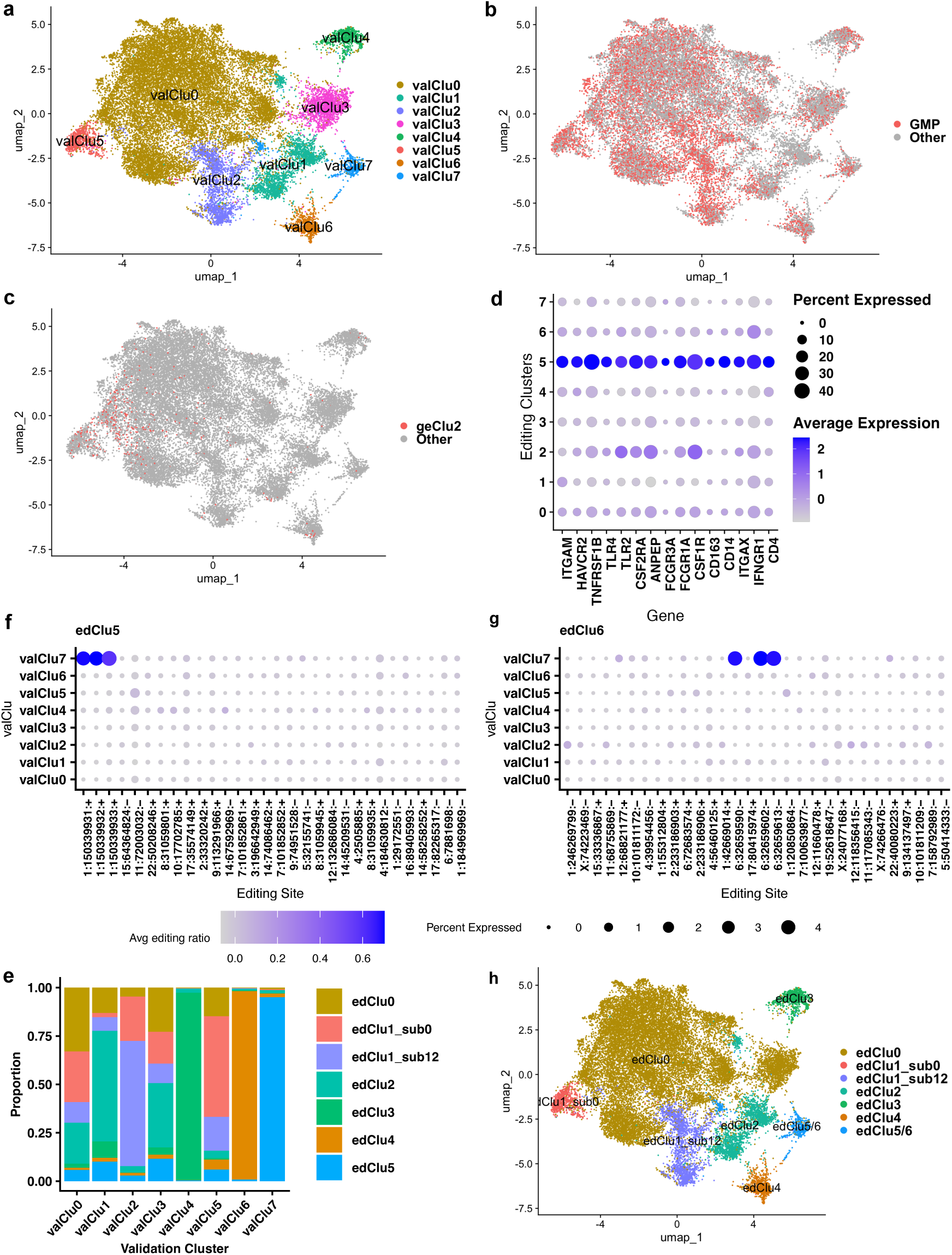
Validation of RNA editing-defined clusters. (a) RNA editing-defined clusters identified from the independent validation cohort. (b) Distribution of GMP cell type. (c) Cells assigned to geClu2 as reported in Ferrall-Fairbanks *et al.* (d) Expression of geClu2-associated marker genes across the eight RNA editing-defined clusters in the validation cohort. (e) Proportional distribution of predicted discovery editing clusters within each validation cluster. (f,g) Average editing ratio of marker editing sites from discovery clusters edClu5 and edClu6 across validation clusters. (h) Final correspondence and assignment between discovery and validation RNA editing-defined clusters.

Despite the lower frequency of mono-biased patients, cells belonging to the previously described adverse transcriptional cluster geClu2 were primarily mapped to valClu5 (**Fig. 6c**) and geClu2 marker genes were significantly enriched in this cluster (**Fig. 6d**), recapitulating the aggressive GMP-associated post-transcriptional program observed in the discovery cohort.

To directly assess alignment between validation clusters and RNA editing-defined clusters from the discovery cohort, we performed reference mapping using Seurat. Each validation cell was assigned to its closest discovery editing cluster, and the results were visualized using UMAP projections and stacked bar plots (**Sup. Fig. 4c**, **Fig. 6e**). Clear alignment patterns emerged, including mapping of valClu2 to edClu1_sub12and valClu5 to edClu1_sub0.

These mappings were further validated by examining the average editing levels of discovery cluster–specific marker editing sites within each validation cluster (**Sup. Tab. 21-22**). Marker editing sites displayed higher editing levels and broader cellular representation in their corresponding validation clusters—particularly valClu2 (edClu1_sub12), valClu4 (edClu3), valClu5 (edClu1_sub0), valClu6 (edClu4), and valClu7 (edClu5/6)—supporting the robustness of the mapping (**Sup. Fig. 5a-5h**). Because valClu7 contained marker sites from both edClu5 and edClu6, it was annotated as edClu5/6 (**Fig. 6f-6g**). The final correspondence between discovery and validation clusters is summarized in **Fig. 6h**.

We next assessed whether clinical and oncogenic associations observed in the discovery cohort were reproducible in the validation dataset. Consistent with discovery results, HMA treatment was associated with increased cellular proportions and MELs in edClu2, edClu4, and edClu5/6, with edClu3 showing a modest increase (**Fig. 7a-7b**). When comparing WHO CMML subtypes using untreated samples, edClu3 and edClu5/6 were again more abundant in CMML-0 patients than in CMML-1/2 patients (**Fig. 7c-7d**), mirroring discovery cohort findings.

**Fig. 7.**
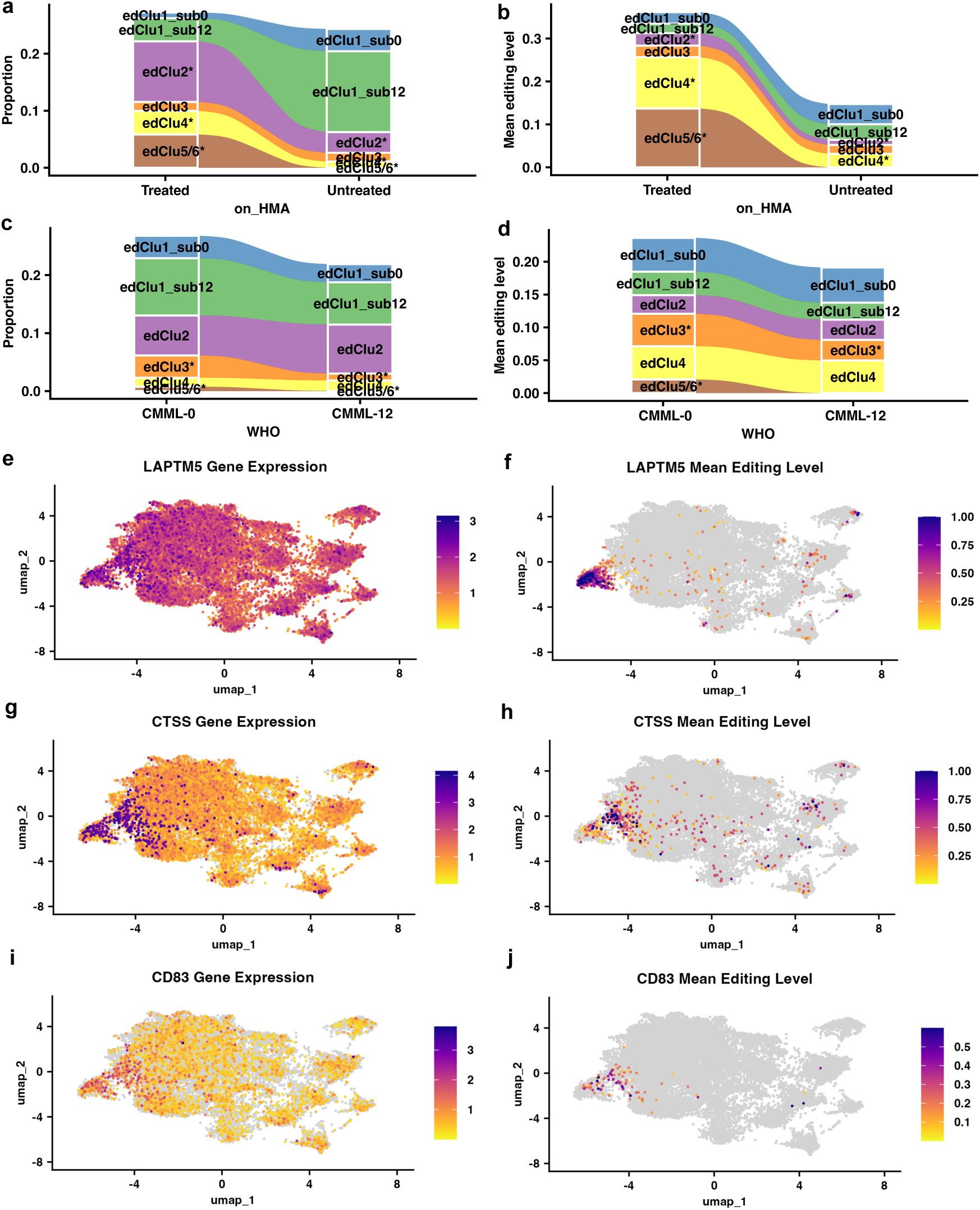
Validation of changes in RNA editing-defined clusters across clinical features and oncogenic editing sites. (a,c) Association between clinical features and the proportion of cells across editing-defined clusters. For each sample, the proportion of cells assigned to each editing cluster was calculated as the number of cells in the cluster divided by the total number of cells in the sample. (b,d) Association between MEL and clinical features. MEL was computed for each editing site by averaging the editing ratios of all cells within each editing cluster and sample. These values were then averaged across all marker editing sites for each cluster. (e,g,i) Gene expression profile of the gene of interest. (f,h,j) Mean editing levels of the gene of interest. * denotes clusters exhibiting apparent changes.

To confirm reproducibility of oncogenic RNA editing programs, we examined high-confidence oncogenic editing sites identified in the discovery cohort. All previously prioritized sites within *ACSL1*, *ADGRE5*, *CD83*, *CTDSP1*, *CTSS*, and *LAPTM5* were also detected in the validation dataset. Among these, *LAPTM5*, *CTSS*, and *CD83* exhibited elevated gene expression specifically within edClu1_sub0, accompanied by significantly increased RNA editing levels at the corresponding sites (**Fig. 7e-7j**). This confirms the significance of the coordinated expression-editing patterns observed in the discovery cohort.

Finally, although the most aggressive editing-defined state, edClu1_sub0, already aligned with the previously reported gene expression–defined cluster geClu2, we further evaluated its clinical relevance. Focusing on monocyte-biased samples, we compared both cellular proportion and MEL between CMML-0 and CMML-1/2 patients, as well as between *TET2*-mutant and non-*TET2*-mutant cases. In both comparisons, edClu1_sub0 exhibited increased cellular proportion and MEL in CMML-1/2 and *TET2*-mutant patients, further supporting the aggressive clinical phenotype associated with this editing-defined state (**Sup. Fig. 6**).

Collectively, these results demonstrate the reproducibility of RNA editing-defined cellular states, their clinical associations, and their linkage to oncogenic immune-related gene regulation in CMML.

### Cluster- and cell-type-specific regulation of RNA editing by *ADAR1* and *ADAR2*

*ADAR1* and *ADAR2* are the primary enzymes responsible for A-to-I RNA editing in mammals. To investigate their roles in shaping RNA editing programs in CMML, we first examined pseudo-bulk expressions of *ADAR1* and *ADAR2* across the 24 discovery samples. An inverse correlation between the two enzymes was observed (**Sup. Fig. 7a**). Because a major isoform of ADAR1 is interferon-inducible[3], we curated a panel of 127 interferon-related and interferon-stimulated genes (ISGs) (Sup. Tab. 23) and incorporated the first two principal components (PCs) derived from this gene set into the correlation analysis. ADAR1 expression positively correlated with the first PC, whereas ADAR2 expression correlated with the second PC (**Sup. Fig. 7a**), suggesting distinct regulatory contexts. However, correlations between sample-level mean editing ratios and ADAR expression did not reach significance (**Sup. Fig. 7b-7c**), indicating that global RNA editing activity is not solely explained by bulk ADAR expression alone.

We next assessed site-specific regulation by performing binomial regression analyses at individual RNA editing sites, modeling pseudo-bulk editing ratios as a function of *ADAR1* and *ADAR2* expression. To account for interferon signaling, we included the first two PCs as covariates. All models were additionally adjusted for age and sex. This analysis identified 830 editing sites significantly associated with *ADAR1* expression and 616 associated with *ADAR2* expression, with 228 sites overlapping between the two enzymes (**Sup. Fig. 7d; Sup. Tab. 24**), supporting site-specific and partially overlapping regulatory roles.

To identify cluster-specific ADAR-associated editing programs, we intersected ADAR-associated sites with cluster-specific marker editing sites (**Sup. Fig. 7e**). *ADAR1*-mediated editing was most pronounced in edClu1_sub0, edClu3, and edClu6, which harbored the largest numbers of *ADAR1*-associated editing sites. *ADAR2*-associated editing was also observed across clusters but generally at lower levels, consistent with a dominant role for *ADAR1* in CMML RNA editing.

We then examined ADAR expression at single-cell resolution across RNA editing-defined clusters. *ADAR1* was broadly expressed across CMML cells, with the highest expression observed in edClu1_sub0, an editing-defined state associated with aggressive clinical features (**Fig. 8a**). In contrast, *ADAR2* expression was comparatively lower in edClu1_sub0, despite being broadly expressed across other clusters (**Fig. 8b**). This high *ADAR1*/low *ADAR2* expression pattern in edClu1_sub0 was independently confirmed in the validation cohort (**Sup. Fig. 8a-8b**).

**Fig. 8.**
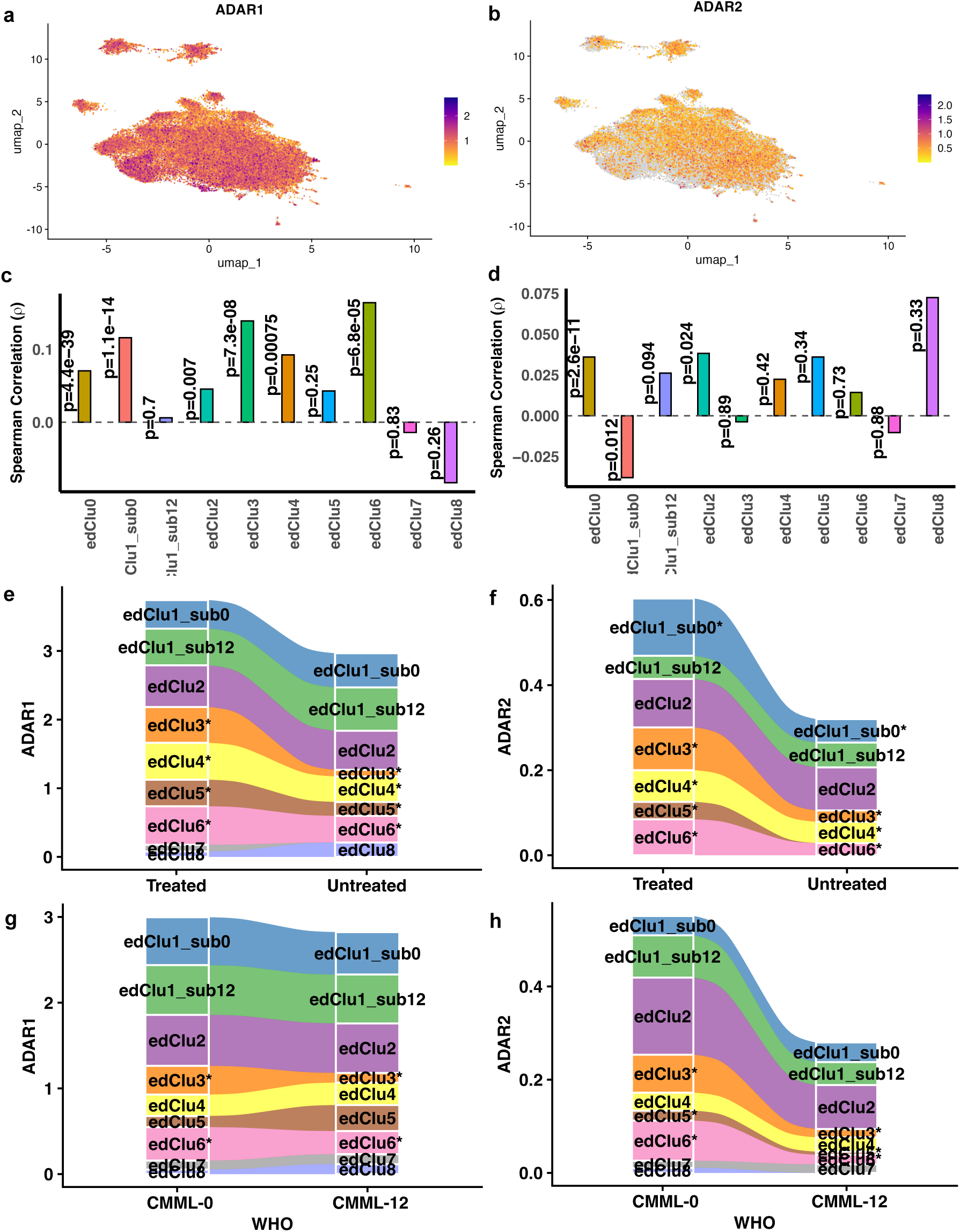
*ADAR1* and *ADAR2* expression patterns and their association with RNA editing across RNA editing-defined clusters. (a, b) Distribution of *ADAR1* and *ADAR2* expression at single-cell level. (c, d) Spearman correlation between *ADAR1* or *ADAR2* expression and MEL across editing-defined clusters. MEL is calculated across all editing sites. (e,f,g,h) *ADAR1* and *ADAR2* expression across RNA editing-defined clusters under different clinical conditions. * denotes clusters exhibiting apparent changes.

To directly assess the relationship between ADAR expression and RNA editing activity at single-cell resolution, we evaluated spearman correlations between *ADAR1* or *ADAR2* expression and MELs within each editing-defined cluster. In edClu1_sub0, edClu3, and edClu6, *ADAR1* expression showed a strong positive correlation with editing activity, whereas *ADAR2* expression was weakly or negatively correlated (**Fig. 8c-8d**). In most other clusters, both *ADAR1* and *ADAR2* showed positive associations with MELs, although the effect was consistently stronger for *ADAR1*. These findings support *ADAR1* as the primary driver of RNA editing in CMML and highlight cluster-specific divergence between *ADAR1* and *ADAR2* regulation.

Finally, we examined *ADAR1* and *ADAR2* expression in relation to clinical features using analyses analogous to those applied to RNA editing clusters. Following HMA treatment, *ADAR1* expression increased in edClu3, edClu4, edClu5, and edClu6 but remained largely unchanged in edClu1_sub0, consistent with observed changes in RNA editing activity. Across WHO CMML subtypes, *ADAR1* expression increased in edClu3 and edClu6 but decreased in edClu5, whereas *ADAR2* expression increased in edClu3, edClu4, edClu5, and edClu6 (**Fig. 8e-8f**). In survival analyses, *ADAR1*-based models identified edClu3 and edClu6 as significantly associated with overall survival (**Sup. Tab. 25**), while *ADAR2*-based models identified edClu3 and edClu7 (**Sup. Tab. 26**). These results indicate that edClu3 and edClu6 represent relatively protective RNA editing states, supported by concordant patterns in RNA editing activity and ADAR expression.

## Discussion

Aberrant A-to-I RNA editing represents an underexplored layer of molecular heterogeneity and regulation in CMML. By developing a single-cell aware RNA editing framework and applying it to independent discovery and validation cohorts, we identified reproducible RNA editing-defined cellular states that can be resolved independently of gene expression and exhibit distinct biological and clinical implications. Notably, a GMP-like editing state (edClu1_sub0) aligned with a previously described inflammatory, monocytic-biased transcriptional program associated with adverse outcomes. Other editing-defined states—particularly edClu3 and edClu6—were enriched in earlier-stage disease, depleted in *TET2*-mutant cases, and associated with improved survival. Additionally, multiple editing-defined states showed systematic increases following HMA treatment. Together, these findings establish RNA editing as a complementary organizational layer that captures clinically meaningful cellular heterogeneity beyond conventional gene expression and mutation-based stratification.

The regulatory relationship between RNA editing enzymes and editing-defined states provides a mechanistic framework that links RNA editing to clinical phenotypes. Site-level modeling revealed extensive but partially non-overlapping sets of *ADAR1*- and *ADAR2*-associated editing sites, supporting locus-specific regulation rather than a uniform global effect. Notably, edClu1_sub0— the editing-defined state most strongly associated with adverse clinical features—consistently exhibited high *ADAR1* and low *ADAR2* expression in both discovery and validation cohorts. Strikingly, the independently derived geClu2 transcriptional state maps onto the same cellular population defined here by RNA editing. These findings support a unifying model in which differential ADAR activity shapes post-transcriptional regulation at sentinel loci and contributes to the inflammatory, monocytic-biased cellular program associated with poor prognosis in CMML.

The identification of convergent increases in both RNA editing levels and expression of *LAPTM5*, *CTSS*, and *CD83* within edClu1_sub0 further illustrates how RNA editing may rewire immune-associated regulatory circuits in aggressive disease states. These genes play central but distinct roles in antigen presentation, immune activation, and endolysosomal function. *LAPTM5* modulates lysosomal trafficking and receptor degradation in hematopoietic cells, *CTSS* provides the protease activity required for invariant-chain cleavage and extracellular matrix remodeling, and *CD83* stabilizes MHC-II and *CD86* by preventing their ubiquitination while regulating T-cell costimulation through membrane-bound and soluble isoforms. Their coordinated upregulation and editing suggest a concerted post-transcriptional mechanism amplifying immune and inflammatory signaling in CMML.

Notably, many prioritized editing events fall within introns and 3′UTRs, regions that influence splicing fidelity, miRNA engagement, and RNA-binding protein occupancy. Editing at these sites may promote isoform switching or post-transcriptional de-repression that amplify these functions, thereby reinforcing inflammatory antigen-presenting programs. Coupled with dominant *ADAR1* activity in edClu1_sub0, these findings support a model in which enzyme-driven remodeling of post-transcriptional regulation contributes to the emergence and maintenance of adverse cellular states. These hypotheses are directly testable using isoform-resolved long-read RNA sequencing, RNA–protein interaction assays, and targeted perturbation of ADAR enzymes or specific editing sites.

This study has several limitations that inform future investigations. The sample size, particularly within specific clinical or genetic strata, limits power for fine-grained genotype editing interaction analyses and for rarer editing states (e.g. like edClu7 and edClu8). The unstranded nature of the scRNA-seq data necessitated stringent strand-aware filtering. Although extensive benchmarking and QC mitigate false positives, orthogonal validation at key loci remains important. Finally, while we show strong associations between enzyme levels, editing states, and clinical outcomes, including the link between edClu1_sub0 and geClu2, establishing causality will require targeted perturbational studies.

## Conclusions

In summary, our study demonstrates that RNA editing defines reproducible cellular states in CMML with distinct biological, clinical, and prognostic relevance. These findings establish RNA editing as a previously underappreciated regulatory layer that is largely independent of conventional gene expression–based clustering and provides new insight into the molecular organization and heterogeneity of CMML. More broadly, this work highlights the potential of single-cell RNA editing analysis to reveal biologically meaningful cellular states in cancer and other complex diseases.

## Methods

### Single cell RNA-seq data

The single cell (sc) RNA-seq FASTQ files for 24 CMML samples were downloaded from GEO: accession number GSE211033, and 3 healthy samples were retrieved from GSE133181. The demographic and clinical characteristics of the patients are provided in **Sup. Tab. 1**.

### Discovery of RNA editing sites from scRNA-seq data

#### Two-phase Discovery of RNA Editing Sites and DNA Mutations

To leverage high sequencing depth at the whole-sample level, we performed the discovery phase using more stringent thresholds. Quantification of the identified RNA editing sites and DNA mutations can then be conducted at the sc level with lower thresholds to enhance sensitivity. This approach balances accuracy and sensitivity, maximizing the opportunity to identify more variants with high precision.

#### Reference-based Detection of Cell Barcodes

We compared the sequenced cell barcode sequences with the reference barcodes, correcting those while tolerating 1 mismatch. This approach not only improves the accuracy of cell barcode assignment but also maximizes the retention of usable reads.

#### Augment the Alignment Strategy

We improved read alignment in two complementary ways. 1) We built on our previous work by using a splice-junction database that enumerates all exon–exon combinations for each annotated human gene (GRCh38). Reads were aligned with Bowtie v1.2.3[16] to both the splice-junction database and the reference genome. We then reconciled results with a stringent selection procedure: when a read mapped uniquely to the same locus in both references, that alignment was retained; when mappings disagreed, we kept the higher-quality alignment based on alignment score and discarded both alignments if the alignment score was equal between the two. 2) Additionally, we performed an independent alignment with STAR v 2.7.9a[17] (two-mode option) and compared its outputs with Bowtie. Prior studies have shown that combining aligners can reduce false positives by ∼50%. Accordingly, for each read we selected the higher-scoring alignment between STAR and Bowtie, and we also retained uniquely aligned reads from either aligner to increase sensitivity for downstream analyses.

#### Remove PCR Duplicates

In most applications, PCR duplicates were commonly removed at the whole-sample level, which can significantly reduce the number of usable reads at the single-cell level. To address this, we checked the cell identity of each read and remove the duplicates that only occur within the same cell.

#### Apply Multiple Filters

We applied stringent filters to minimize technical and genomic artifacts. 1) Genetic polymorphisms: removed sites overlapping with the dbSNP build 151 SNVs and NCBI “common” human variants (https://www.ncbi.nlm.nih.gov/variation/docs/human_variation_vcf/. 2) Problematic regions: excluded loci within simple repeats (UCSC simpleRepeat v10.3.28) and rRNA annotated by the UCSC RepeatMasker track. 3) Strand bias: used a two-sided Fisher’s exact test and sites with Benjamini–Hochberg (BH) FDR < 0.05 were discarded. 4) Positional bias: applied a distance-to-read-start test and sites with BH FDR < 0.05 were discarded. 5) Nearby mutations: removed sites with any non–A>G/T>C variant within ±150 bp (read length). 6) Recurrence/coverage: retained sites observed in ≥10 samples.

#### Annotation of RNA editing sites

RNA editing sites were annotated to GRCh38 using annovar v498[29]. Due to the nature of this study, the RNA sequencing is not stranded. We corrected A->G and C->T variants based on the strand of their host reference gene. Therefore, a portion of T->C may be A->G if they arise from the complementary strand of reference genome and if they located in non-annotated reference gene region.

#### Quantification of Single-cell RNA-editing

From the uniquely aligned, duplicate-removed BAM, we split reads by cell barcode to generate per-cell BAMs using scripts from SComatic[30]. For each discovered editing site in each cell, we counted edited and unedited reads, computed total coverage, and calculated the editing ratio (edited/total). Sites detected in at least 10 cells were retained for downstream analyses.

### Quality Control, Clustering and Cell Type Assignment

The RNA editing ratio data were then integrated into a Seurat[31] object using the R package (version 5.2.1). Cells with zero editing ratios across all editing sites were excluded from further analysis. Additionally, editing sites with more than 90% of their values equal to 1.00 across all cells were removed. The remaining data were then scaled and principal component analysis (PCA) was performed for linear dimensionality reduction. The top 20 principal components were selected based on an elbow plot for downstream analysis. To account for batch effects, the Harmony algorithm[32] was applied to the PCA-reduced data. Cell clustering was conducted using Seurat’s Louvain algorithm with a resolution parameter set to 0.07, resulting in the identification of nine editing-based clusters labeled edClu0 through edClu8. To further investigate the heterogeneity within edClu1, we performed sub clustering at the same resolution (0.07), identifying three subclusters. Of these, subcluster zero was retained and labeled as edClu1_sub0, while the other two subclusters were merged and labeled edClu1_sub12. This process resulted in a final set of ten editing clusters, consisting of the original eight clusters plus two subclusters derived from edClu1. Non-linear dimensionality reduction was performed using Uniform Manifold Approximation and Projection (UMAP) to visualize the editing clusters. Unless otherwise specified, all Seurat functions were performed using default parameters. Cell type annotations based on gene expression were assigned by matching cell barcodes from the gene expression dataset published in Ferrall-Fairbanks *et al* (2022)[18] to the corresponding barcodes in the editing dataset.

### Mapping Gene Expression to Editing Data

To incorporate gene expression data with RNA editing data, we first matched cell barcodes and identified the intersection of cells shared between the gene expression and editing datasets. Gene expression counts for common cells were extracted from the gene expression data provided in Ferrall-Fairbanks *et al*.[18] Gene expression counts were integrated as a separate assay for a Seurat object. Genes with no expression across all cells were removed and the data were then normalized using Seurat’s log normalization method. Then, only these common cells were used for downstream analyses which involves single-cell gene expression.

### Differential Expression Analysis of Editing Clusters

To identify differentially expressed RNA editing sites between distinct editing-defined clusters and cell types, we used the FindAllMarkers() function from Seurat. This applies the Wilcoxon Rank Sum test by default. Editing sites with an adjusted p-value < 0.05 and a log₂ fold change > 1 were considered significant and retained as differential markers for downstream analysis.

### Calculation of Mean RNA Editing at Single Cell Level

For each cell, the overall mean RNA editing level was calculated by averaging the editing ratios across all detected sites within that cell. Additionally, for each cell, a cluster-specific mean editing level was calculated by averaging the editing ratios of the marker editing sites associated with its corresponding cluster.

### Calculation of Average Gene Expression Across Clusters

To estimate average gene expression levels across clusters, we calculated the mean expression for each gene using log-normalized single-cell RNA-seq data. Log-normalized expression values were first transformed back to the linear scale using the expm1() function in R. The mean expression was then computed on this linear scale across all cells within each cluster. Finally, the resulting mean values were re-transformed to the log scale using the log1p() function to ensure consistency with other log-transformed data used in the analysis.

### Differential Expression Analysis of Cell Type Marker Editing Sites

To identify cell type-based RNA editing markers, we first annotated cells with reference cell type identities based on gene expression data. Using these annotations, we grouped cells by cell type and performed differential analysis on the editing ratios to identify editing sites that were specifically enriched in each cell type. Editing sites with an adjusted p-value < 0.05 and a log₂ fold change > 1 were considered significant and retained as differential markers for downstream analysis.

### Functional Enrichment Analysis

Functional enrichment analysis was performed using g:Profiler[33] with the human reference genome (GRCh38). Genes harboring marker editing sites with p-value < 0.05 and a log₂ fold change > 1 was input into g:Profiler to identify enriched biological processes and pathways.

### Summary of the Biological Functions Grouped by Editing Cluster

To provide a clear and interpretable summary of the biological functions, enriched terms were grouped by editing cluster. The top-ranked terms (based on statistical significance and biological relevance) were then categorized into major biological function themes. Results were shown in Sup. Fig. 2j.

This categorization was guided by keyword-based mapping and performed using ChatGPT-4.0 (OpenAI, 2025) with the following query:

*“Group the functions by cluster. Then, categorize the top enriched terms for each cluster into major biological function categories (e.g., ‘Immune response and signaling’ and ‘DNA repair & chromatin remodeling’, etc.). Under each category, list key representative terms or pathways in a concise summary format (2–5 categories per cluster). Draw a table with first column is editing cluster and second column major categories using bullet points and associated functions.”*

### Computation of Sample and Cluster-Level Editing Metrices to Find the Association of Editing Clusters with Clinical Features

For each editing cluster and sample, we calculated two metrics: the proportion of cells in the cluster and the mean editing level (MEL). The proportion was defined as the number of cells in a given cluster within a given sample, divided by the total number of cells in that sample. For each editing site, we calculated the MEL for each cluster within each sample by averaging the editing ratios of all cells in that cluster. This yielded a matrix for each editing site with samples as rows and clusters as columns. To obtain a summary measure, we then averaged these values across all marker editing sites for each cluster, resulting in a single matrix that represents the average MEL per cluster per sample. To ensure consistency, we included all editing clusters across all 24 CMML samples, imputing zeros for clusters not present in a given sample.

### Association with Clinical Features

To explore clinical associations with proportions and MEL of each cluster, we filtered samples based on clinical annotations. For comparisons involving bias classification (as defined by Ferrall-Fairbanks *et al*., 2022) and World Health Organization (WHO) classification subtypes, we excluded five samples treated with hypomethylating agents (HMA) and used 19 untreated samples. In the WHO classification comparisons, CMML-1 and CMML-2 were combined into a single group. For HMA-treated vs. untreated comparisons, we focused on 10 “Normal-like” bias samples, excluding one sample (SRR21016551) due to inconsistent clinical annotation. FAB classification (MDS-CMML vs. MPN-CMML) was analyzed using 23 samples (excluding one with missing data). For mutation-based comparisons, samples were grouped into TET2-only and no mutation, using 11 samples with available mutation data (9 samples had missing information and rest are TET2_SRSF2).

### Logistic Regression for CMML Classification

To evaluate the potential of RNA editing levels as biomarkers for CMML classification, we applied logistic regression models separately to each editing cluster. The binary outcome variable was CMML subtype, coded as 1 for CMML-12 (combined CMML-1 and CMML-2) and 0 for CMML-0. The predictor variable was the mean RNA editing level of each cluster for a given sample.

For cluster 𝑐, the model was specified as

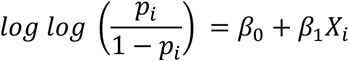

where:

- 𝑝_𝑖_ is the probability that sample 𝑖 belongs to the CMML-12 subtype,
- 𝑋_𝑖_ is the mean RNA editing level of the selected cluster for sample 𝑖,
- 𝛽_0_ and 𝛽_1_ are model parameters estimated from the data.

Model fitting was performed using the glm() function in R with a binomial family and logit link. For each cluster, the model was compared to a null model (intercept-only) using a likelihood ratio test (LRT) to assess the statistical significance of the editing cluster as a predictor.

Predicted probabilities from each fitted model were used to classify samples with a threshold of 0.5. Model performance was evaluated by computing accuracy, sensitivity, and specificity based on the resulting confusion matrix.

Additionally, receiver operating characteristic (ROC) curve analysis was performed using the ROCR package in R. The area under the ROC curve (AUC) was calculated to quantify the discriminatory ability of each model. ROC curves were generated by plotting true positive rates against false positive rates across all classification thresholds.

### CNV-Based Stratification of Healthy-like and Cancer-like Cells Using inferCNV

To distinguish cells associated with malignant hematopoiesis from those resembling normal hematopoiesis at single-cell resolution, we performed copy number variation (CNV)–based stratification using inferCNV (https://github.com/broadinstitute/inferCNV). The analysis was conducted on the gene expression matrix derived from 24 CMML samples together with three healthy donor samples used as references. The input dataset comprised 27,076 genes and 101,131 single cells (82,014 CMML cells and 19,117 healthy reference cells).

InferCNV was run with a cutoff threshold of 0.1, retaining 8,468 genes for downstream analysis. The algorithm generated a CNV-normalized expression matrix in which each value reflects the deviation of gene expression from a neutral baseline. For each cell, a CNV score was computed as the mean squared deviation of expression values across retained genes. Cells with CNV scores at or below the 10th percentile were classified as healthy-like, reflecting minimal copy number alterations. Cells at or above the 80th percentile were classified as cancer-like, indicative of pronounced CNV gains or losses. Using these thresholds, 16,403 cancer-like cells and 8,202 healthy-like cells were identified in the gene expression dataset. After mapping CNV labels to the RNA editing dataset, 7,873 cancer-like cells and 6,325 healthy-like cells were retained for downstream analyses.

### Differential Analysis of RNA Editing Between Healthy-like and Cancer-like cells

To identify RNA editing sites that were differentially edited between healthy-like and cancer-like cells at single-cell resolution, we used the Wilcoxon rank-sum test implemented in the FindMarkers() function in Seurat. Editing sites with adjusted *P* values < 0.05 were considered significant. Sites with log₂ fold changes > 1 were classified as upregulated in cancer-like cells, whereas sites with log₂ fold changes < −1 were classified as upregulated in healthy-like cells (i.e., downregulated in cancer-like cells).

### Interferon-stimulated Gene Sets

We curated interferon-stimulated genes (ISGs) from the literature[34–37] and included only those that were also present in the gene expression data of common cells with matched RNA editing information. The following ISG sets were incorporated in the analysis: 13 IFN-α and IFN-γ response genes from the HUGO Gene Nomenclature Committee, 52 IU-dsRNA-responsive genes, 38 ISG RS genes, 39 ISGliu genes, and 21 ISGchronic genes, as reported in previous studies.

### Psuedo-bulk Gene Expression Matrix

Pseudo-bulk gene expression profiles were constructed by aggregating raw counts across all cells within each of the 24 samples, capturing expression for 26,701 genes. Counts per million (CPM) were then computed and genes with CPM > 1 in at least 10 samples were retained, resulting in 12,878 genes for downstream analysis. A pseudo count of 0.01 was added to the CPM values, followed by log₁₀ transformation and scaling.

### Covariate Construction for Binomial Regression

To account for potential regulatory influences on editing, a set of 108 ISGs present in the filtered pseudo-bulk gene expression dataset were selected. PCA was performed on these ISGs, with the first two principal components explaining approximately 50% of the total variance (PC1: 32.90%, PC2: 17.37%). The final covariate matrix used in the regression analyses included scaled pseudo-bulk expression values of ADAR and ADARB1, along with PC1 and PC2 derived from ISG expression.

### Association between RNA editing and Myelomonocytic Proliferation

To assess the relationship between RNA editing and myelomonocytic proliferation, we performed binomial regression analyses at bulk level. For each RNA editing site, the number of edited reads in sample 𝑖 was modeled as

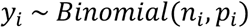

where:

- 𝑦_𝑖_ is the number of edited reads at a specific RNA editing site for the *i*-th sample,
- 𝑛_𝑖_ is the total number of reads covering that site, and
- 𝑝_𝑖_ is the underlying editing ratio (i.e., probability of success).

The log-odds of 𝑝_𝑖_ were modeled using a logistic function:

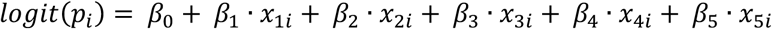

Where:

- 𝑥_1_ is monocyte absolute number or monocyte percentage
- 𝑥_2_ and 𝑥_3_ is the scaled pseudo-bulk expression of ADAR1 and ADAR2
- 𝑥_4_ is age
- 𝑥_5_ is gender

Models were fitted using the glm() function in R with a binomial family and logit link. Bulk edited-read and total-read matrices initially contained 509,543 sites across 24 samples. We restricted the analysis to sites also detected in the single-cell dataset (3,326 sites) and further retained 2,354 sites with non-zero edited read counts in at least 10 samples. We applied the BH procedure to account for multiple testing across RNA editing sites and those with adjusted p-values < 0.05 were considered statistically significant.

### Identification of Oncogenic and Anti-oncogenic RNA Editing Sites

To identify RNA editing sites with potential oncogenic relevance, we applied an integrative approach based on the overlap of three complementary strategies:

1. Editing Cluster-Based Selection (edClu1_sub0): We first selected significantly upregulated RNA editing sites from the edClu1_sub0 editing cluster, which is closely related to GMP-like cells. Sites were filtered based on an adjusted p-value < 0.05 and log₂ fold change >1.
2. Disease-State Enrichment (Cancer-like): We next identified editing sites upregulated in cancer-like cells using inferCNV-based differential editing analysis at the single-cell level. Sites meeting the criteria of adjusted p-value < 0.05 and log₂ fold change > 1 were retained. These were combined with editing sites uniquely enriched in cancer-like cells.
3. Association with Myeloid Disease Features: Finally, we selected editing sites that were positively associated with either absolute number of monocytes or monocytes%, based on regression models adjusting for key covariates. Only sites with positive effect sizes and adjusted p-values < 0.05 in either models were retained.

### Survival Analysis using Cox Proportional Hazards Models

To assess the prognostic relevance of RNA editing activity, we fit Cox proportional hazards models separately within each editing cluster, restricting the analysis to patients who had not received HMAs. For each cluster, we considered four models with the following predictors:

1. the proportion of cells,
2. the mean RNA editing level,
3. *ADAR1* gene expression, and
4. *ADAR2* gene expression.

All models were also adjusted for age and sex. Survival time and vital status for each patient were obtained from the clinical phenotype dataset.

The Cox model used for each cluster was:

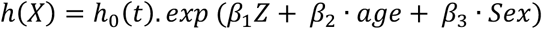

where ℎ(𝑋) is the hazard at time 𝑡, ℎ_0_(𝑡) is the beaseline hazard, 𝑍 represents the cluster-specific predictor of interest. Hazard ratios (HR), 95% confidence intervals (CI), and p-values were extracted for covariate analysis to assess the association with survival. The editing clusters were ranked by p-value significance to highlight clusters with the most prognostic relevance. Analysis was carried out for the common cells between editing and gene expression.

### Validation Samples

An independent set of 16 CMML samples from 15 patients (including 2 samples from a single patient) was used for validation. Raw FASTQ files were downloaded from the Gene Expression Omnibus (GEO) under accession number GSE211033. Reads were processed using the same preprocessing workflow and RNA editing detection pipeline as applied to the discovery dataset. Three samples (SRR21016553, SRR21016555, and SRR21016556) with fewer than 50 detected editing sites were excluded from downstream analysis.

The remaining samples were integrated into a single Seurat object. Editing ratios for a curated set of 1,308 high-confidence RNA editing sites were used as input features for clustering analysis. The resulting editing ratio matrix was analyzed using the same Seurat pipeline and parameters as in the discovery dataset, including clustering at resolution RNA_snn_res.0.07.

### Mapping of Validation Clusters to Discovery-defined Editing Clusters

To enable comparison between editing-defined clusters in the discovery and validation datasets, we employed Seurat’s label transfer workflow. First, FindTransferAnchors() was used to identify shared feature space between the discovery (reference) and validation (query) datasets based on the editing ratio matrices. These anchors were then used with TransferData() to assign each cell in the validation dataset a predicted identity corresponding to one of the editing clusters in the discovery set. To further validate this correspondence, we examined the average editing ratio of marker editing sites in the discovery set within each validation cluster. For each marker editing site, we computed the average editing ratio and the percentage of cells with detectable editing.

### Computation of Sample and Cluster-level Editing Metrices for Validation Data

To ensure consistency with the discovery dataset, we computed the same two metrics in the validation data to assess clinical relevance: the proportion of cells and the MEL per editing cluster and per sample. However, to calculate MELs in the validation set, we restricted the analysis to a set of common marker editing sites shared between discovery and validation clusters.

These common markers were identified by performing differential editing analysis within the validation dataset using the transferred cluster annotations. Marker sites were defined as those with an adjusted p-value < 0.05 and log_2_ fold change > 1. We then intersected these validation-derived markers with the original marker sites from the discovery dataset to obtain a shared set of informative editing sites. MELs were computed by averaging editing ratios across these shared markers for each cluster within each sample, resulting in a cluster-by-sample validation matrix comparable to the discovery set.

### Bulk mean RNA Editing Calculation

Bulk-level RNA editing was assessed using the original bulk editing count and total read matrices that included 509,543 editing sites across 24 samples. These sites were filtered to include only those also detected in the single-cell dataset, reducing the set to 3,326 sites. An additional filtering step retained 2,354 sites with non-zero editing counts in at least 10 samples. For each sample, the mean bulk editing level was calculated as the average editing ratio across all retained sites.

### Binomial Regression Analysis of RNA Editing and ADAR Expression at Bulk Level

To assess the relationship between RNA editing and ADAR expression, we applied the same binomial regression framework described above for myelomonocytic proliferation. We included the scaled pseudo-bulk expression of *ADAR1*, the first two principal components (PC1 and PC2) derived from ISG expression, age, and sex as predictors. The same analysis was repeated using *ADAR2*.

### Correlation between *ADAR1* and *ADAR2* Expression and mean RNA Editing Ratios Across Different Editing Clusters

For each editing-defined cluster, we computed Pearson correlation coefficients between the single cell expression levels of *ADAR1* and *ADAR2* and the mean editing ratios calculate across all editing sites, using only the cells belonging to that specific cluster.

## List of abbreviations

3′UTR: 3′ untranslated region
ADAR: Adenosine deaminase acting on RNA
A-to-I: Adenosine-to-inosine
AUCs: Area under the operating characteristic curve
BH: Benjamini–Hochberg
CMML: Chronic myelomonocytic leukemia
CNV: Copy number variation
C-to-U: Cytidine-to-uridine
GMP: Granulocyte–monocyte progenitor
GO: Gene Ontology
GO:BP: Gene Ontology biological process
HMA: Hypomethylating agent
HSCs: Hematopoietic stem cells
ISGs: Interferon-related and interferon-stimulated genes
MELs: Mean editing levels
MEP: Erythroid progenitor
scRNA-seq: Single-cell RNA sequencing
UMAP: Uniform Manifold Approximation and Projection

## Declarations

### Ethics approval and consent to participate

All data was de-identified and derived from publicly available datasets. No new patient data was presented in this body of work.

### Consent for publication

Not applicable.

### Availability of data and materials

The scRNA-seq data and associated clinical annotations analyzed in this study are publicly available through the Gene Expression Omnibus (GEO) with GEO accession number: GSE211033. And the three healthy individuals are available with GEO submission number: GSE133181.

The code used for sc data analysis is available at https://github.com/tjgu/CMML-scRNAediting/tree/main.

### Authors’ contributions

TG conceived and designed this study and performed RNA editing sites discovery. NW performed the single-cell RNA editing analysis. TG and NW drafted the manuscript. DB maintained and refined the RNA editing discovery pipeline. SN and EP provided the dataset. SN, MF, EP and MW contributed critical discussions. All authors reviewed, edited and approved the final version of the manuscript.

## Acknowledgements

We thank our colleagues, Drs. Subramaniam Malarkannan, Demin Wang, and Renren Wen (Versiti Blood Research Institute), for their thoughtful discussions and suggestions. We thank Emily Diaz, PhD, for assistance with the preparation of the manuscript.

