## Supplementary Figures for "Single-cell RNA editing defines clinically relevant cellular states in chronic myelomonocytic leukemia"

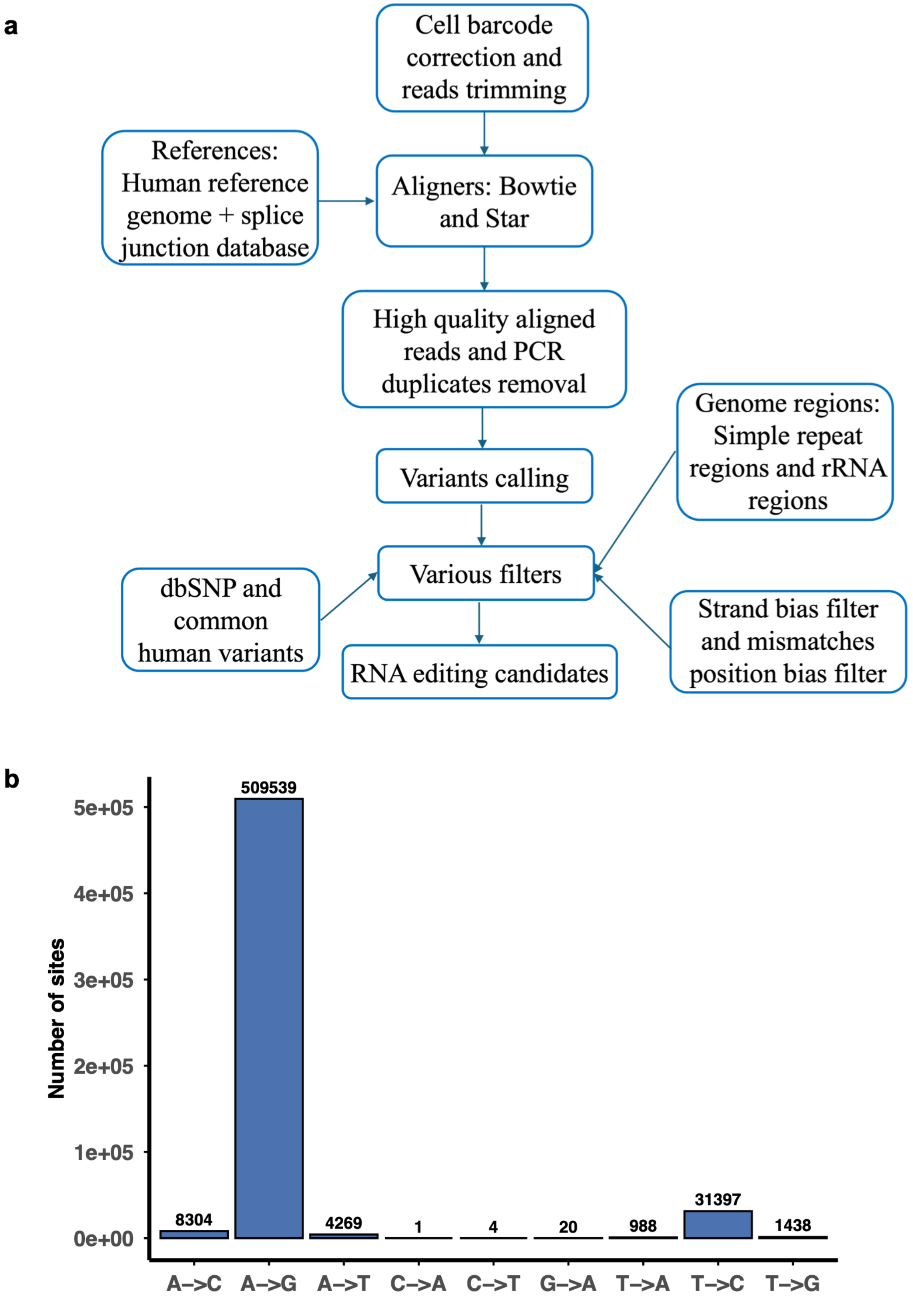


**Sup. Fig. 1. RNA editing site discovery pipeline and the distribution of mutation types.** (a) RNA editing sites detection pipeline. (b) The distribution of all identified variant sites, including potential DNA mutations.

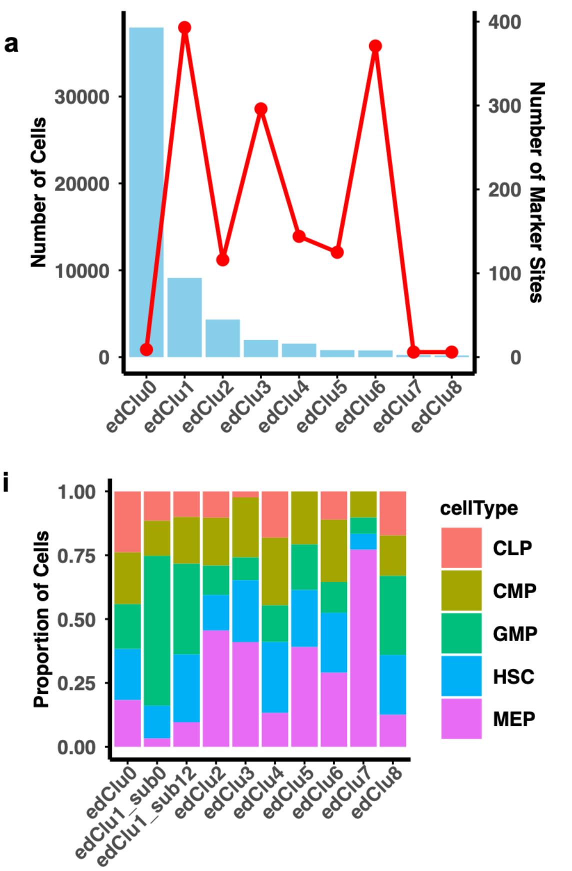


j

| Editing Cluster | Dominant Biological Function Categories & Key Terms |
|---|---|
| edClu1_sub0 | • Immune response & signaling: immune response, signaling, response to stimulus<br>• Signal transduction & cytokine activity: MAPK cascade, phosphorylation<br>• Cell communication & activation: positive regulation of cell communication, immune activation, inflammatory response, cytokines, pathogens |
| edClu1_sub12 | • Phosphorylation & protein modification: peptidyl-serine phosphorylation, protein modification processes<br>• Lymphoid/hematopoietic processes: lymphoid progenitor differentiation, hematopoietic progenitor differentiation<br>• DNA repair & telomere regulation: homologous recombination, telomere maintenance |
| edClu2 | • DNA repair & chromatin remodeling: recombinational repair, chromatin organization<br>• RNA and protein metabolism: regulation of RNA metabolic process, translational initiation<br>• Stress response: cellular response to stress, DNA strand invasion |
| edClu3 | • Translation & RNA processes: cytoplasmic translation, regulation of mRNA metabolic process<br>• Metabolic regulation: gene expression, biosynthetic processes<br>• Implication: Likely less differentiated or stem/progenitor states |
| edClu4 | • Macromolecule metabolism & cell cycle: regulation of biosynthesis, cell cycle<br>• Positive regulation of gene expression: transcription control, biosynthesis<br>• Possibly related to myeloid-biased hematopoietic cells |
| edClu5 | • Protein degradation & catabolism: proteolysis, protein catabolic process<br>• Cell cycle & turnover: regulation of cell cycle, apoptotic signaling<br>• Implication: Enriched for proteasome-associated pathways, may reflect proliferative or stress-responsive state |
| edClu6 | • RNA/protein regulation: RNA metabolic process, translation initiation<br>• Cell regulation: leukocyte-mediated immunity, chromatin modification<br>• May reflect specific immune features of activated cell types |

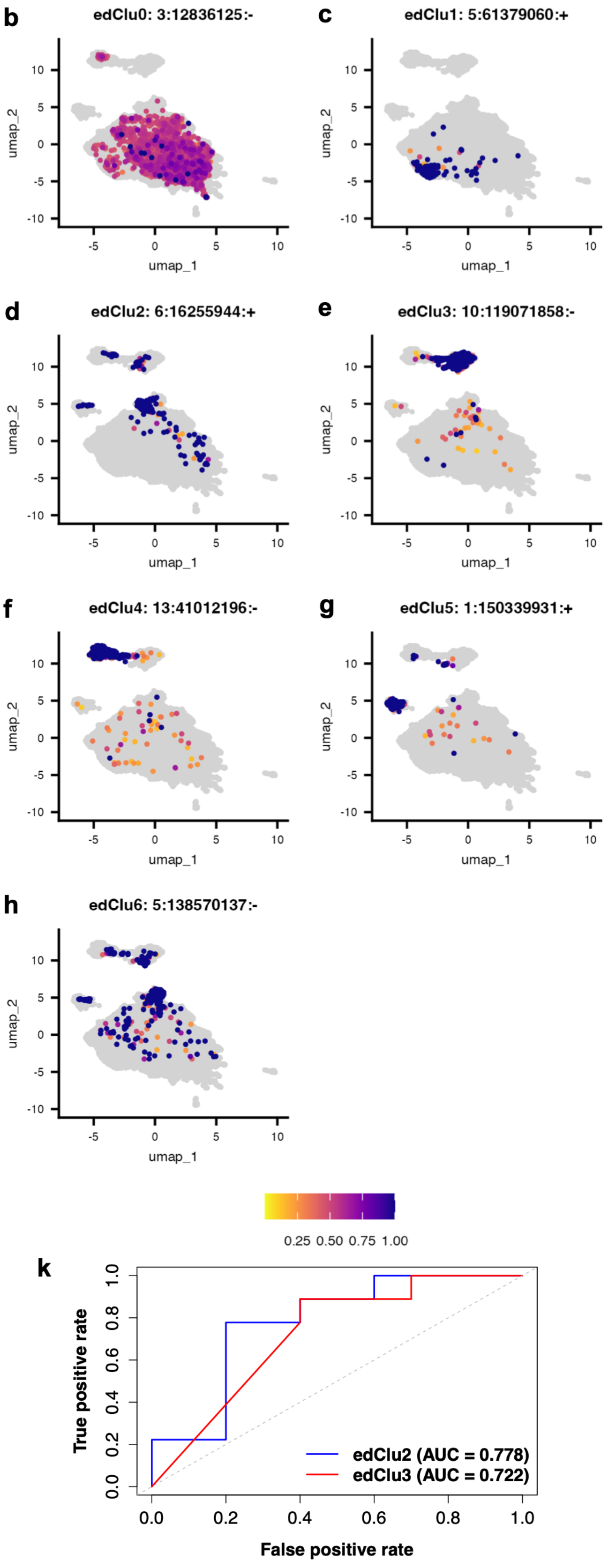


**Sup. Fig. 2. RNA editing clusters and their characteristics.** (a) Number of marker editing sites and cells present in each cluster. (b-h) Distribution of editing ratios for top marker editing sites across their respective editing clusters. (i) Relative distribution of cells from each annotated cell types across editing-defined clusters. (j) ChatGPT-assisted curation for editing clusters. (k) ROC curve for predicting CMML subtype using mean editing levels of each cluster.

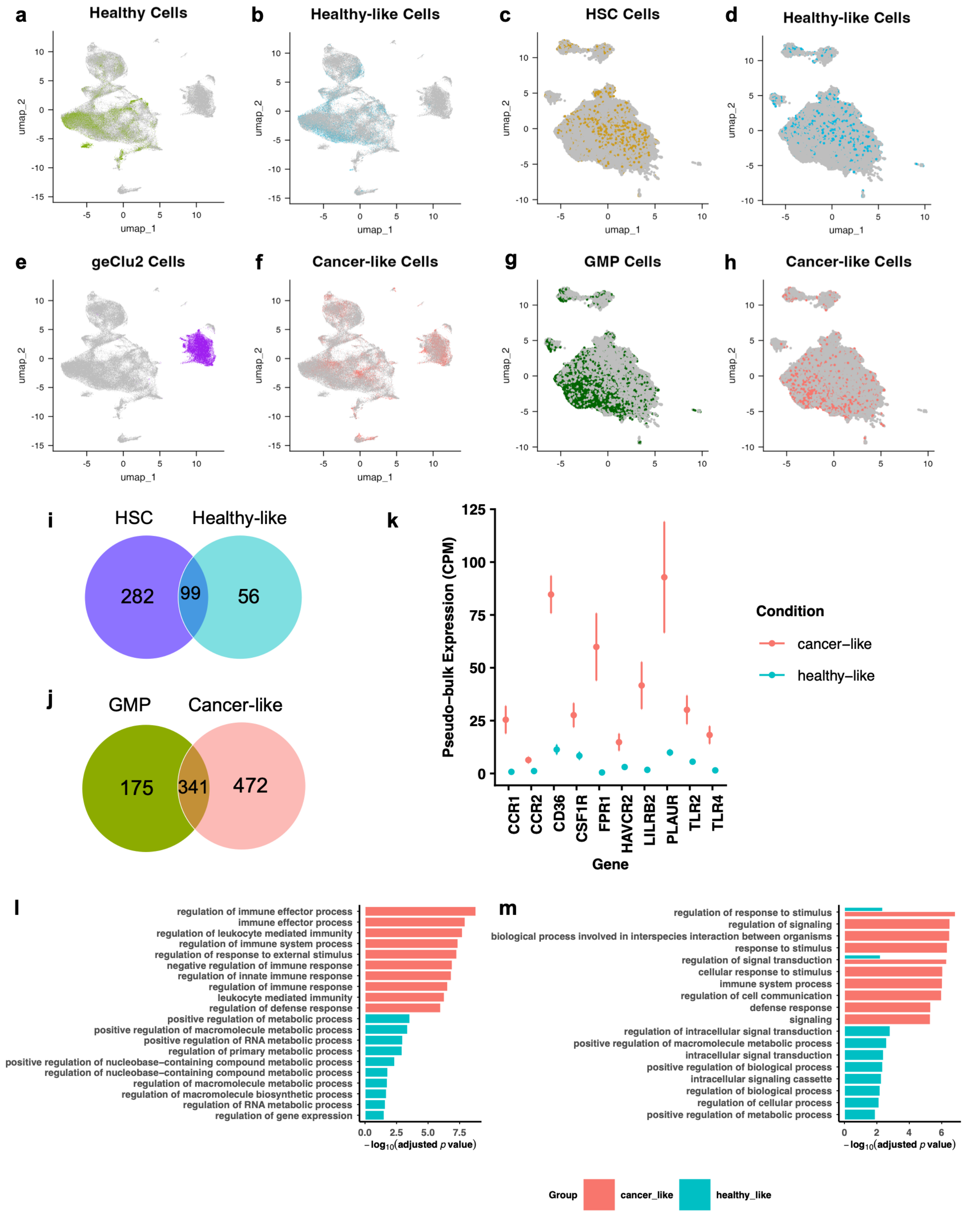


**Sup. Fig. 3. Identification and functional characterization of healthy-like and cancer-like cells**. **(**a,b) Distribution of healthy and healthy-like cells across gene expression-defined clusters. (c, d) Distribution of HSC and healthy-like cells across RNA editing-defined clusters. (e) Gene expression UMAP of geClu2 cells. (f) Gene expression UMAP of cancer-like cells. (g) RNA editing UMAP for GMP cells. (h) RNA editing UMAP for cancer-like cells. (i) Overlap between HSC and healthy-like marker genes. (j) Overlap between GMP and cancer-like marker genes. (k) Differential expression of the established receptors identified from the comparison between healthy and CMML CD34+ cells in cancer-like and healthy-like cell population. (l) Gene functions for genes harboring differential expressed marker editing sites between healthy-like and cancer-like. (m) Functional enrichment analysis for the genes harboring cancer-like or healthy-like unique editing sites.

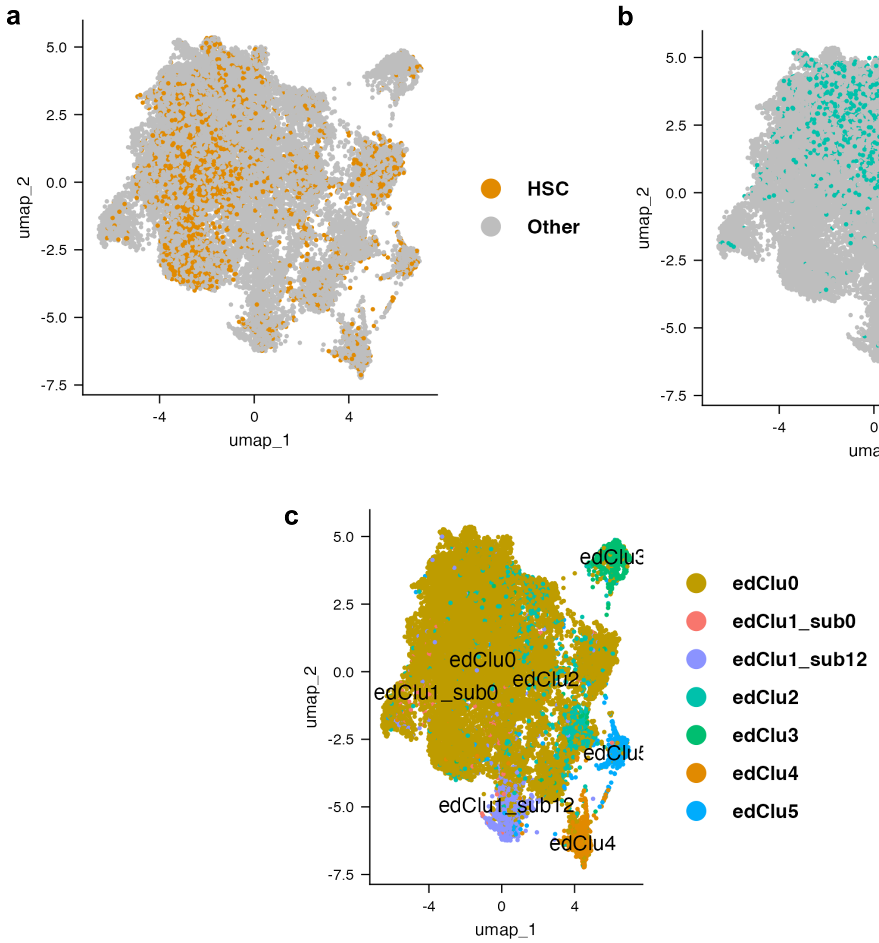


**Sup. Fig. 4. Map cell-types and RNA editing clusters identified from the discovery cohort onto the validation clusters.** (a) Distribution of HSC cell type across validation clusters. (b) Distribution of MEP cell type across validation clusters. (c) Correspondence between validation clusters and the editing-defined clusters from the discovery dataset.

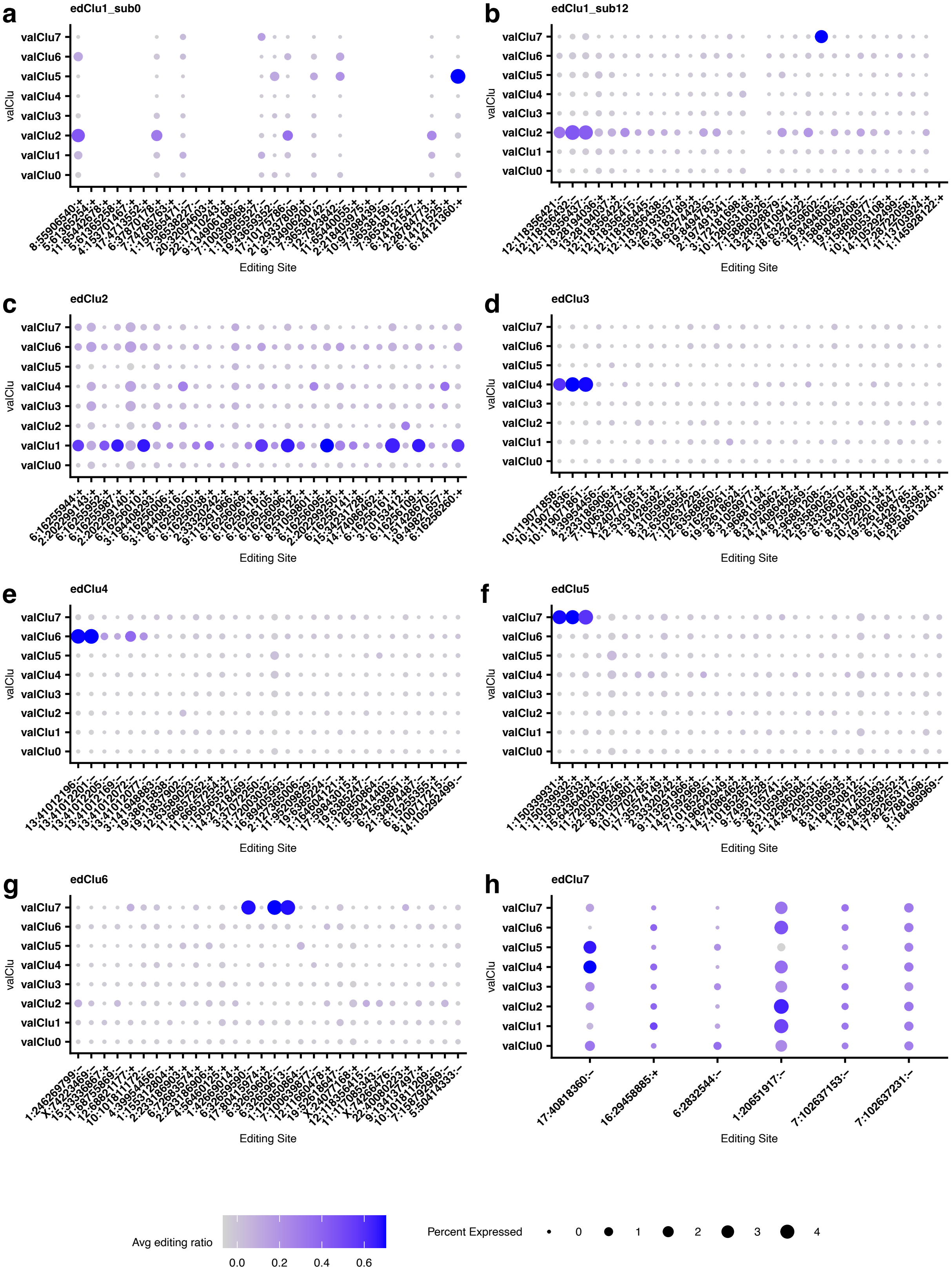


**Sup. Fig. 5. Distribution of average editing ratio of marker editing sites from the discovery cohort in each validation cluster.** (a – h) Average editing ratio of marker editing sites from each discovery cluster within each validation cluster.

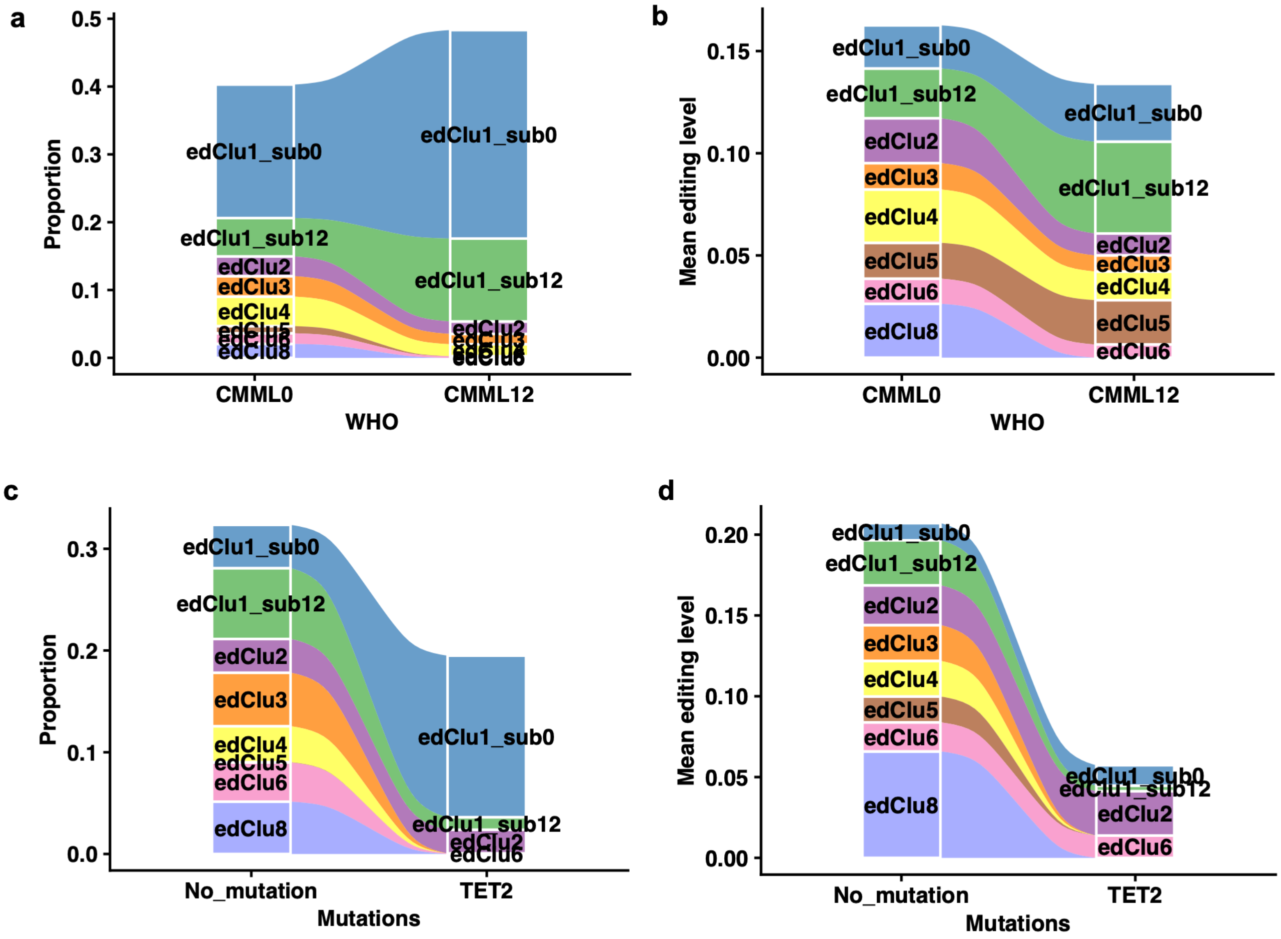


**Sup. Fig. 6**. **Alterations in RNA editing–defined clusters across clinical features**. (a,c) Association between clinical features and the proportion of cells across editing-defined clusters. For each sample, the proportion of cells assigned to each editing cluster was calculated as the number of cells in the cluster divided by the total number of cells in the sample. (b,d) Association between mean editing level (MEL) and clinical features. MEL was calculated for each editing site by averaging the editing ratios of all cells within each editing cluster and sample. These values were then averaged across all marker editing sites for each cluster.

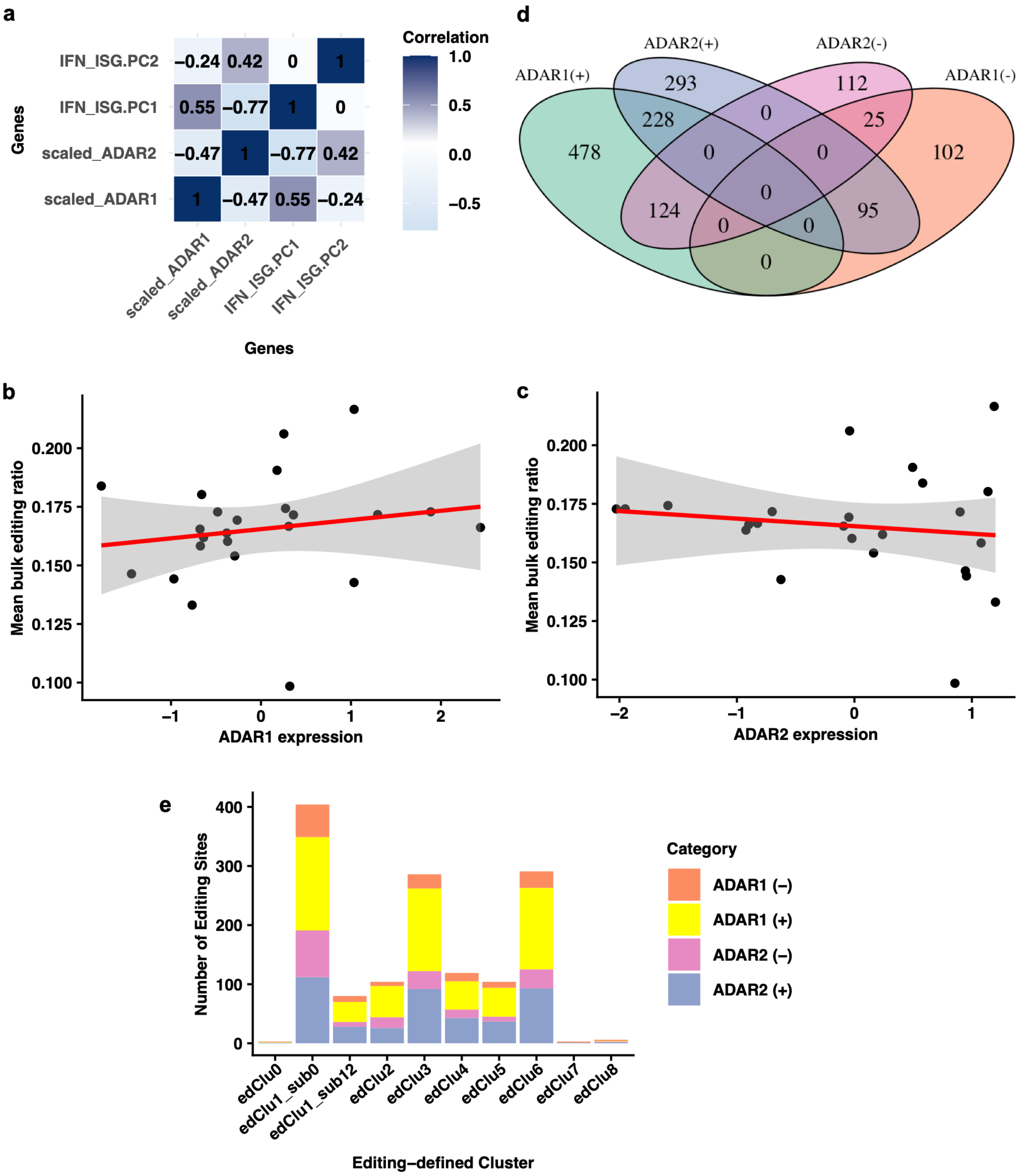


**Sup. Fig. 7. Cluster–specific expression patterns of *ADAR1* and *ADAR2*.** (a) Correlation of *ADAR1* and *ADAR2* expression at bulk level. IFN_ISG_PC1 and IFN_ISG_PC2 represent the top two PCs derived from 127 interferon-related and interferon-stimulated genes (ISGs). (b-c) Correlation of *ADAR1* and *ADAR2* expression with mean bulk editing ratio. (d) Editing sites associated with *ADAR1* and *ADAR2* expression at bulk level. (e) ADAR-associated editing sites in editing clusters at bulk level.

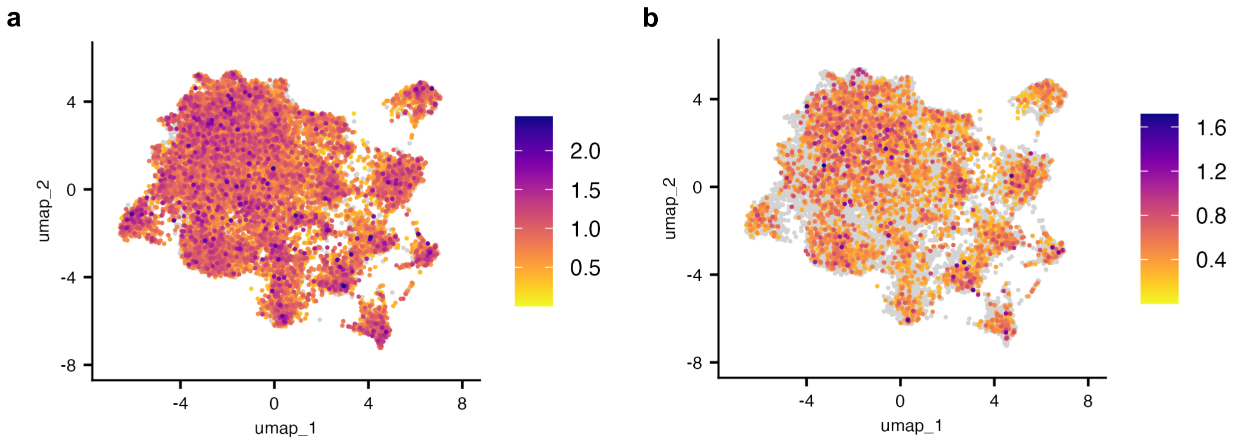


**Sup. Fig. 8. *ADAR1* and *ADAR2* expression across validation samples.** (a) Distribution of *ADAR1* expression across validation samples. (b) Distribution of *ADAR2* expression across validation samples.
